# Nanopore long reads enable the first complete genome assembly of a Malaysian *Vibrio parahaemolyticus* isolate bearing the pVa plasmid associated with acute hepatopancreatic necrosis disease

**DOI:** 10.1101/861476

**Authors:** Han Ming Gan, Christopher M. Austin

**Affiliations:** Centre for Integrative Ecology, School of Life and Environmental Sciences, Deakin University, 3220 Geelong, Victoria, Australia; Deakin Genomics Centre, Deakin University, 3220 Geelong, Victoria, Australia; School of Science, Monash University Malaysia, 47500 Petaling Jaya, Selangor, Malaysia

## Abstract

**Background:** *Vibrio parahaemolyticus* MVP1 was isolated from a Malaysian aquaculture farm affected with shrimp acute hepatopancreatic necrosis disease (AHPND). Its genome was previously sequenced on the Illumina MiSeq platform and assembled *de novo* producing a relatively fragmented assembly. Despite identifying the binary toxin genes in the MVP1 draft genome that were linked to AHPND, the toxin genes were localized on a very small contig precluding proper analysis of gene neighbourhood.

**Methods:** The genome of Vibrio parahaemolyticus MVP1 was sequenced on the Nanopore MinION device to obtain long reads that can span longer repeats and improve genome contiguity. *De novo* genome assembly was subsequently performed using long-read only assembler (Flye) followed by genome polishing as well as hybrid assembler (Unicycler).

**Results:** Long-read only assembly produced three complete circular MVP1 contigs consisting of chromosome 1, chromosome 2 and the pVa plasmid that *pirAB^vp^* binary toxin genes. Polishing of the long read assembly with Illumina short reads was necessary to remove indel errors. The complete assembly of the pVa plasmid could not be achieved using Illumina reads due to the presence of identical repetitive elements flanking the binary toxin genes leading to multiple contigs. Whereas these regions were fully spanned by the Nanopore long reads resulting in a single contig. In addition, alignment of Illumina reads to the complete genome assembly indicated there is sequencing bias as read depth was lowest in low-GC genomic regions. Comparative genomic analysis revealed the presence of a gene cluster coding for additional insecticidal toxins in chromosome 2 of MVP1 that may further contribute to host pathogenesis pending functional validation. Scanning of all publicly available *V. parahaemolyticus* genomes revealed the presence of a single AinS-family quorum-sensing system in this species that can be targeted for future microbial management.

**Conclusions:** We generated the first chromosome-scale genome assembly of a Malaysian *pirAB^Vp^*-bearing *V. parahaemolyticus* isolate. Structural variations identified from comparative genomic analysis provide new insights into the genomic features of *V. parahaemolyticus* MVP1 that may be associated with host colonization and pathogenicity.

## Introduction

*Vibrio parahaemolyticus* is a marine gram-negative *Enterobacteriaceae* that has been recognized as an important pathogen affecting commercially-relevant shrimp species such as the giant tiger prawn (*Penaeus monodon*) and the Pacific white shrimp (*Litopenaeus vannamei*) ^1–3^. In recent years, *V. parahaemolyticus* has been linked to acute hepatopancreatic necrosis disease (AHPND)^4, 5^. An AHPND-causing *V. parahaemolyticus* isolate is thought to produce a homolog of the insecticidal *Photorhabdus* insect-related (Pir) binary toxin that induces prolonged damage to the shrimp tissue^2^. AHPND preferentially affects shrimp post-larvae or juveniles with nearly 100% mortality rate within 30-35 days post-stocking^6^. Studies suggest that AHPND originated from China in 2009 and subsequently spread to South East Asia^7^. In Malaysia, AHPND was first reported in 2011 where it still persists in shrimp farms^8^.

The implementation of effective microbial management in aquaculture requires genomic surveillance data that can provide accurate information regarding the geographical origin, virulence factors and antibiotic resistance of a sequenced microbial strain^9^. In Malaysia, this remains challenging due to the paucity of available genomic resources for the relevant pathogens. To date, a majority of the microbial genomic studies in Malaysia consist of single draft genome reports mostly focusing on clinical pathogens without substantial comparative genomics^10–13^. However, recent years have seen a modest increase in the number of studies with more comprehensive sequencing dataset and analysis^14–16^. For example, Yan *et al* sequenced and assembled the draft genomes of 40 Malaysian *V. parahaemolyticus* isolates associated with shrimp aquaculture and performed comparative genomics of more than Asian 100 *V. parahaemolyticus* genomes from public databases. *In-silico* multi-locus sequencing typing and phylogenomic analyses indicate that several Malaysian *V. parahaemolyticus* isolates belong to previously undescribed sequence types and genomic lineages^17^ and recommended further studies be undertaken.

The *pirAB^vp^* genes coding for AHPND-causing binary toxins are localized on a 70-kb pVa plasmid^2^. Surprisingly, despite the small size of the plasmid relative to the chromosomal genome, a complete *de novo* assembly of the pVa plasmid using Illumina-only short reads has not been achieved to date. Recent studies suggested that this is due to the presence of repetitive elements on the pVa plasmid that are longer than the Illumina read length^18^. As a result, *pirAB*-containing contigs that were assembled from Illumina reads are generally short with limited gene content, precluding detailed analysis of the *pirAB* gene neighborhood and their stability in the plasmidome^19^. *V. parahaemolyticus* isolate MVP1 is one of the first *pirAB^Vp^* -harboring *V. parahaemolyticus* isolated from Malaysia to have its genome sequenced^11^. Isolate MVP1 was previously sequenced on an Illumina MiSeq followed by *de novo* genome assembly using the SPAdes assembler. While the *pirAB^vp^* genes could be identified in the draft genome, the assembly was problematic as these were localized on a short contig flanked by small fragments of a transposase gene.

Recent years have seen the democratization of long-read sequencing enabled by Nanopore technology. In contrast to another popular long-read sequencing platform, PacBio, Nanopore sequencing requires minimal lab footprint and very low capital investment. Importantly, it is the first sequencing technology that allows native sequencing, thus eliminating sequencing bias associated with the activity of *Taq*-polymerase. Long reads have been utilized for *de novo* genome assembly in two distinct ways. First, long reads can be used directly in genome assembly followed by polishing with Illumina short reads to improve consensus accuracy^20^. An alternative approach utilizes long-reads to reorder and link contigs that were initially assembled from Illumina reads^21, 22^. While the final assembly produced with this hybrid approach is highly accurate, the contiguity of the assembly is dependent on the quality of the initial Illumina assembly^23^.

Leveraging on the availability of Illumina dataset for isolate MVP1, we aimed to improve on the original genome assembly^11^ through the addition of 40× Nanopore read coverage. We first performed a Nanopore-led assembly using the long-read Flye assembler^24^ followed by polishing with Illumina reads. In addition, we generated a hybrid assembly using Unicycler assembler^23^. Although both methods produced a complete circular pVa plasmid sequence with intact *pirAB^vp^*, complete chromosome assembly was only achieved with the Nanopore-led Flye assembly. This superior assembly provides new insight into the genomic features of *V. parahaemolyticus* MVP1 and will allow a greater understanding of host pathogenicity through comparative genomic analysis with selected *V. parahaemolyticus* isolates.

## Methods

### Nanopore sequencing

Sample collection, gDNA isolation and Illumina sequencing of *V. parahaemolyticus* isolate MVP1 have been previously reported^11^. For Nanopore sequencing, 1 µg of unfragmented gDNA was processed using the now obsolete Nanopore SQK-NSK007 library preparation kit. Sequencing was subsequently performed on an R9 flowcell attached to a MinION device for 48 hours. Base-calling of the produced raw fast5 files used Guppy v3.1 (high accuracy mode) that only is available to Oxford Nanopore Technology (ONT) customers via the ONT community site (https://community.nanoporetech.com).

### De novo genome assembly and genome polishing

Illumina-only assembly and hybrid assembly incorporating high-quality (q>7) Nanopore long reads were performed using Unicycler v.0.4.7^23^. Only contigs equal or larger than 1,000 bp were retained for subsequent analyses. A long-read only *de novo* assembly was also performed with Flye v2.4.2 utilizing similar Nanopore sequencing data^24^. Illumina polishing of the Flye assembly used the unicycler polish tool in Unicycler v.0.4.7 that performed multiple rounds of bwa-mem-based Illumina read alignment to the assembly followed by polishing with Pilon v1.22^25^. Genome completeness was assessed with BUSCO v3 (Gammaproteobacteria odb9) based on the whole proteome from each genome assembly that was predicted using Prodigal v 2.6.3 (default setting) ^26, 27^.

### Comparative Genomic analysis and protein homology modeling

Automated annotation of the whole-genome was carried out using the NCBI Prokaryotic Genome Annotation Pipeline (PGAP)^28^. The translated coding sequences generated from PGAP were subject to additional functional annotation using <u>InterProScan</u>. Intra-chromosomal structural variations and sequence similarity were visualized with BLAST ring image generator (BRIG) using the blastN (e-value = 0.001 cutoff). Sequencing depth was also calculated in BRIG based on the BAM alignment file generated from the bwa-mem mapping of Illumina paired-end reads to the whole-genome^29^. Structural modeling of the putative insecticidal toxin proteins identified in MVP1 used the online Phyre2 webserver (http://www.sbg.bio.ic.ac.uk/phyre2)^30^.

### Identification and phylogenetic analysis of LuxM, OpaM and AinS proteins

Vibrio proteins containing the InterPro signature IPR035304, corresponding to the AinS-family N-acyl-homoserine-lactone autoinducer synthase, were downloaded from the UniProt database ^31^ on the 7^th^ of November 2019 and clustered at 98% amino acid identity cutoff using cd-hit v4.6 ^32^. The protein clusters were subsequently aligned with MAFFT v7.31 ^33^ (”--maxiterate 1000 ‒localpair” setting) followed by maximum likelihood tree construction using FastTree v2.1 ^34^. Calculation and visualization of pairwise protein sequence divergence used SDT v1.2 ^35^. In addition, to confirm the absence of the LuxI-type autoinducer synthase homolog in the *Vibrio parahaemolyticus* proteomes, additional domain search specifically for PF00765 (LuxI autoinducer synthase domain) was performed followed by subsequent InterProScan validation as previously described ^36^.

## Results and discussion

### Chromosomal assembly enabled by Nanopore long reads

A total of 231 megabases of Nanopore data contained in 30,315 reads (N50 = 10,908 bp) were produced from a single Nanopore MINION run on the discontinued R9 flowcell. The base-called Nanopore fastq file has been deposited in the SRA database under the accession code <u>SRX6759854</u>. An initial Unicycler assembly using previously generated Illumina reads (<u>SRX6759853</u>) yielded a relatively fragmented genome (198 contigs; N50 length = 54,109 bp). Supplementing the assembler with approximately 40 x coverage of Nanopore long reads led a substantial improvement in the Unicycler assembly contiguity (9 contigs; N50 length = 1,738,848 bp). Of the 9 assembled contigs, one of them was flagged as “complete” with a contig length of 70kb. On the other hand, the Flye assembly using only Nanopore reads produced three contigs all flagged as “complete and circular”. Despite generating the most contiguous assembly, the initial Flye assembly has the worst BUSCO score with 37.4% BUSCO single-copy and 45.1% fragmented BUSCO genes (Table 1). Polishing of the Flye assembly with Illumina paired-end reads restored the genome completeness to a level that is comparable to the hybrid and Illumina-only assemblies (Table 1 and *Extended data*: Supplemental File 1). The high percentage of fragmented genes in long-read-only assembly is usually a result of erroneous frameshifts from indel sequencing errors in the long reads^20^. Although higher coverage of long read may lead to an increase the consensus accuracy, its accuracy will unlikely match that of an Illumina-polished assembly due to Nanopore-specific systematic errors. As a result, polishing of long-read assembly using Illumina short reads can be considered as a cost-effective but less convenient approach for producing highly accurate and contiguous microbial genome assembly.

**Table 1.**
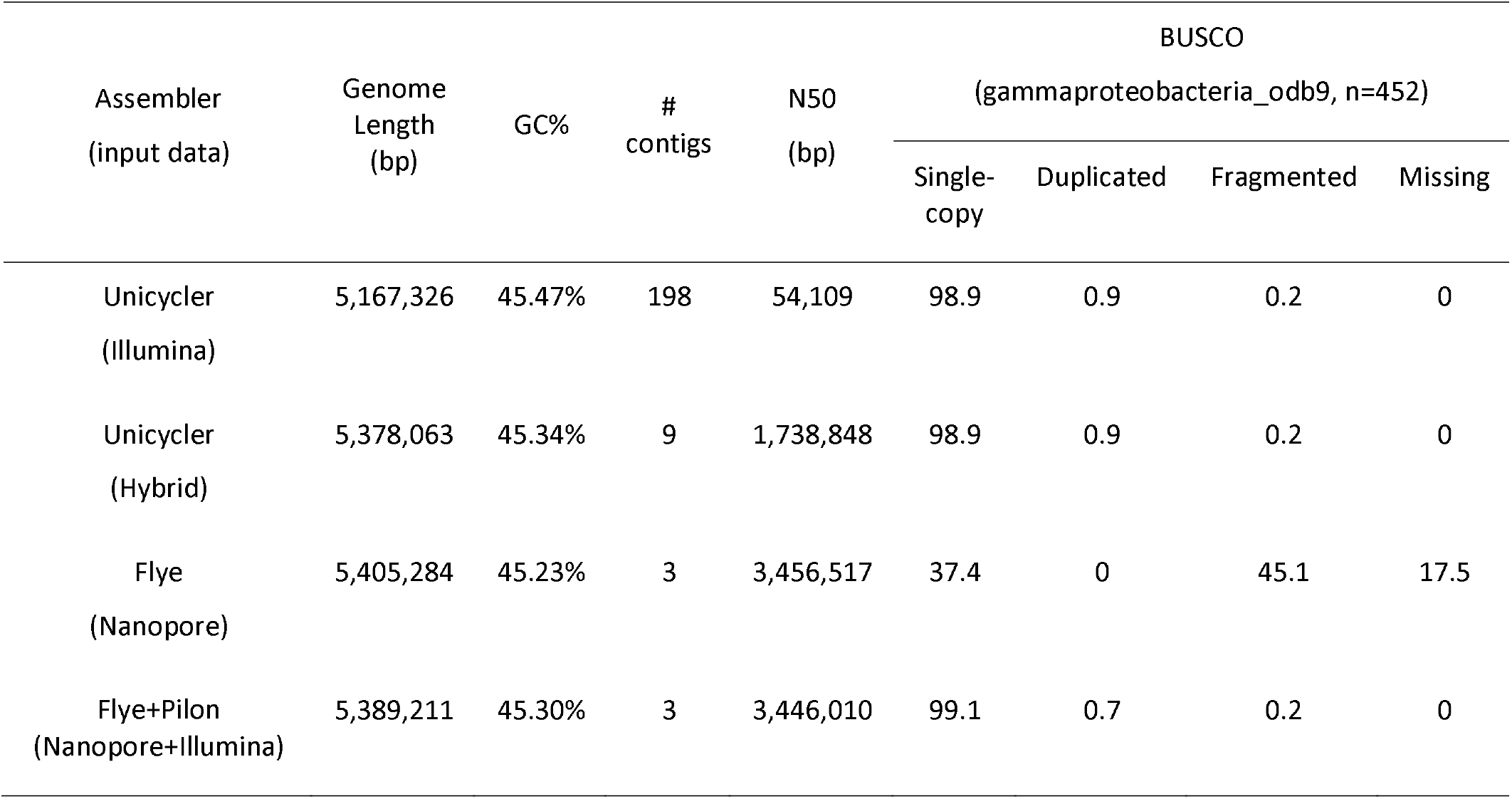
Statistics and BUSCO completeness assessment of the *Vibrio parahaemolyticus* MVP1 assembly.

| Assembler<br>(input data) | Genome<br>Length<br>(bp) | GC% | #<br>contigs | N50<br>(bp) | BUSCO<br>(gammaproteobacteria_odb9, n=452) |  |  |  |
| --- | --- | --- | --- | --- | --- | --- | --- | --- |
|  |  |  |  |  | Single-copy | Duplicated | Fragmented | Missing |
| Unicycler<br>(Illumina) | 5,167,326 | 45.47% | 198 | 54,109 | 98.9 | 0.9 | 0.2 | 0 |
| Unicycler<br>(Hybrid) | 5,378,063 | 45.34% | 9 | 1,738,848 | 98.9 | 0.9 | 0.2 | 0 |
| Flye<br>(Nanopore) | 5,405,284 | 45.23% | 3 | 3,456,517 | 37.4 | 0 | 45.1 | 17.5 |
| Flye+Pilon<br>(Nanopore+Illumina) | 5,389,211 | 45.30% | 3 | 3,446,010 | 99.1 | 0.7 | 0.2 | 0 |

### Reduced Illumina sequencing depth in several low-GC genomic regions

The alignment of Illumina reads to the final chromosomal assembly revealed substantial lower sequencing depths in genomic regions with low GC-content (<30%) (Figure 1). Some of these regions are unique to MVP1 and may risk being overlooked in the Illumina-only assembly at lower sequencing coverage. For example, less than 5x sequencing depth was observed in the region 3,240 to 3,280 kb that contain various putative carbohydrate metabolism genes that are unique to isolate MVP1 in this current genomic comparison. The incorporation of PCR during library preparation is known to reduce the representation of high-AT genomic regions due to the polymerase amplification bias towards GC-balanced fragments^37^. Reduced Illumina sequencing depth in high-AT genomic regions affecting biological interpretation has also been reported in several crustacean mitogenomic libraries as both the phylogenetically informative control region and 16S rRNA genes are relatively AT-rich^38, 39^. Depending on the amount of DNA available for sequencing, a few modifications to the Illumina library preparation step can be considered, such as PCR-free preparation for sample with high starting DNA (> 100 ng) and reduced PCR cycle coupled with higher (>100× genome coverage) sequencing depth for samples with low starting DNA.

**Figure 1.**
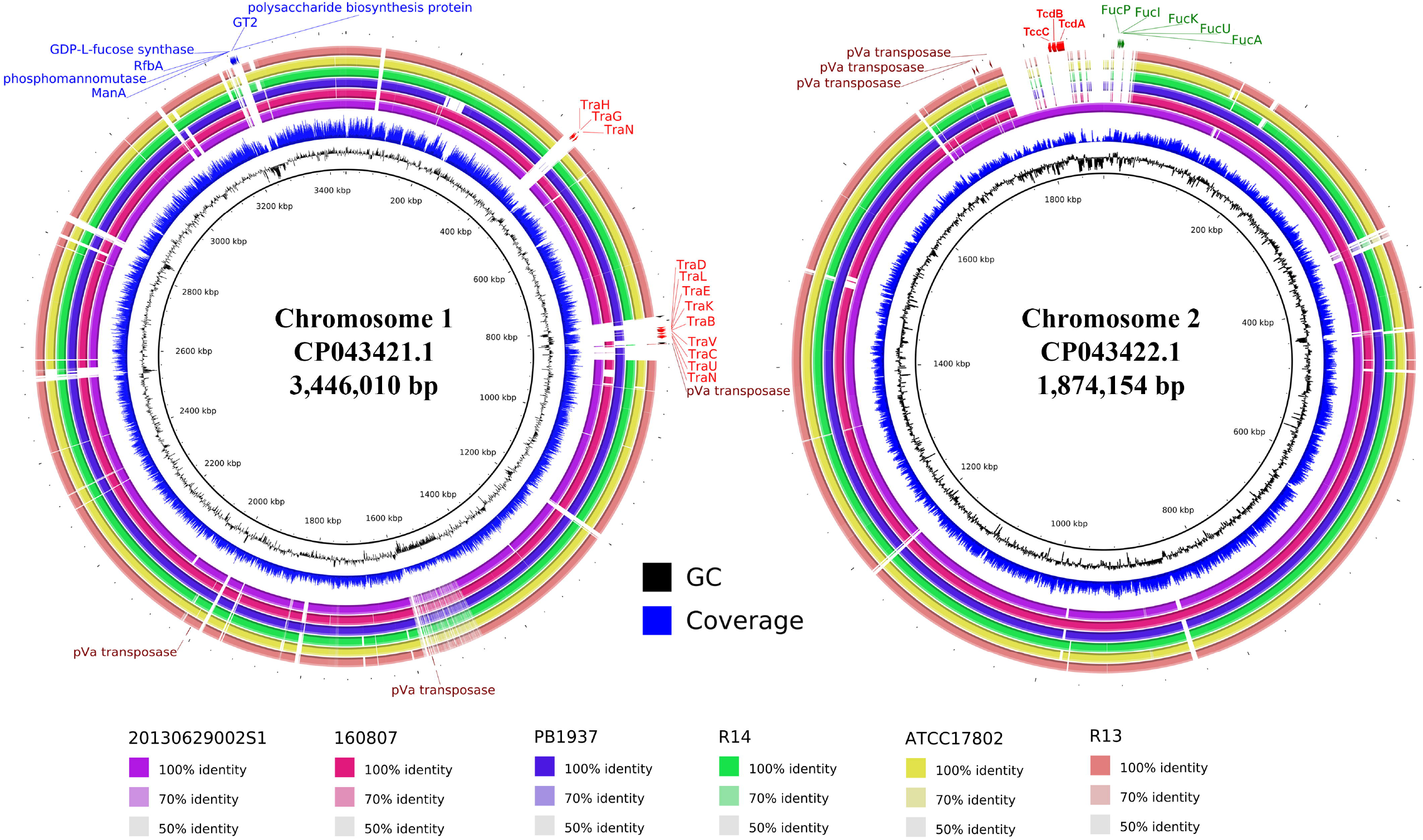
Genomic comparisons of pVa plasmid-harboring *Vibrio parahaemolyticus*. Left, *V. parahaemolyticus* MVP1 chromosome 1 compared against five other *V. parahaemolyticus* chromosome 1. Right, *V. parahaemolyticus* MVP1 chromosome 2 compared against five other *V. parahaemolyticus* chromosome 2. The innermost rings show GC content while the second innermost rings show Illumina sequencing depth along the genome (blue). The remaining rings indicate BLAST comparisons of other V. parahaemolyticus genomes against MVP genome assembly. Arrows indicate genes of interest and were labeled with the predicted protein names.

### Multiple localization of IS5 family transposase genes in the complete genomes of pVa-carrying *V. parahaemolyticus*

In the Illumina-only assembly, the pVa plasmid was assembled into three contigs (Contig24: 63.4 kb, Contig164: 3.4kb; Contig198: 1 kb). Contig198 contains a 921-bp gene that codes for an intact IS5 family transposase and could be aligned equally well to two genomic regions flanking the *pirAB^Vp^* genes in the complete plasmid assembly (Figure 2). This observation can be explained by the inability of Illumina reads to fully span the repeats thus causing them to be collapsed into a single contig. As a result, the localization of *pirAB^Vp^* on the pVa plasmid cannot be confirmed from an Illumina-only assembly. In contrast the entire *pirAB^Vp^* genes and their flanking transposase genes were fully covered by a number of Nanopore long reads enabling the complete assembly of the pVa plasmid in addition to supporting the localization of *pirAB^Vp^* genes on the pVa plasmid. Nanopore read depth compared to Illumina, is relatively even across the aligned genomic region, which is consistent with the absence of PCR bias in the Nanopore library preparation.

**Figure 2.**
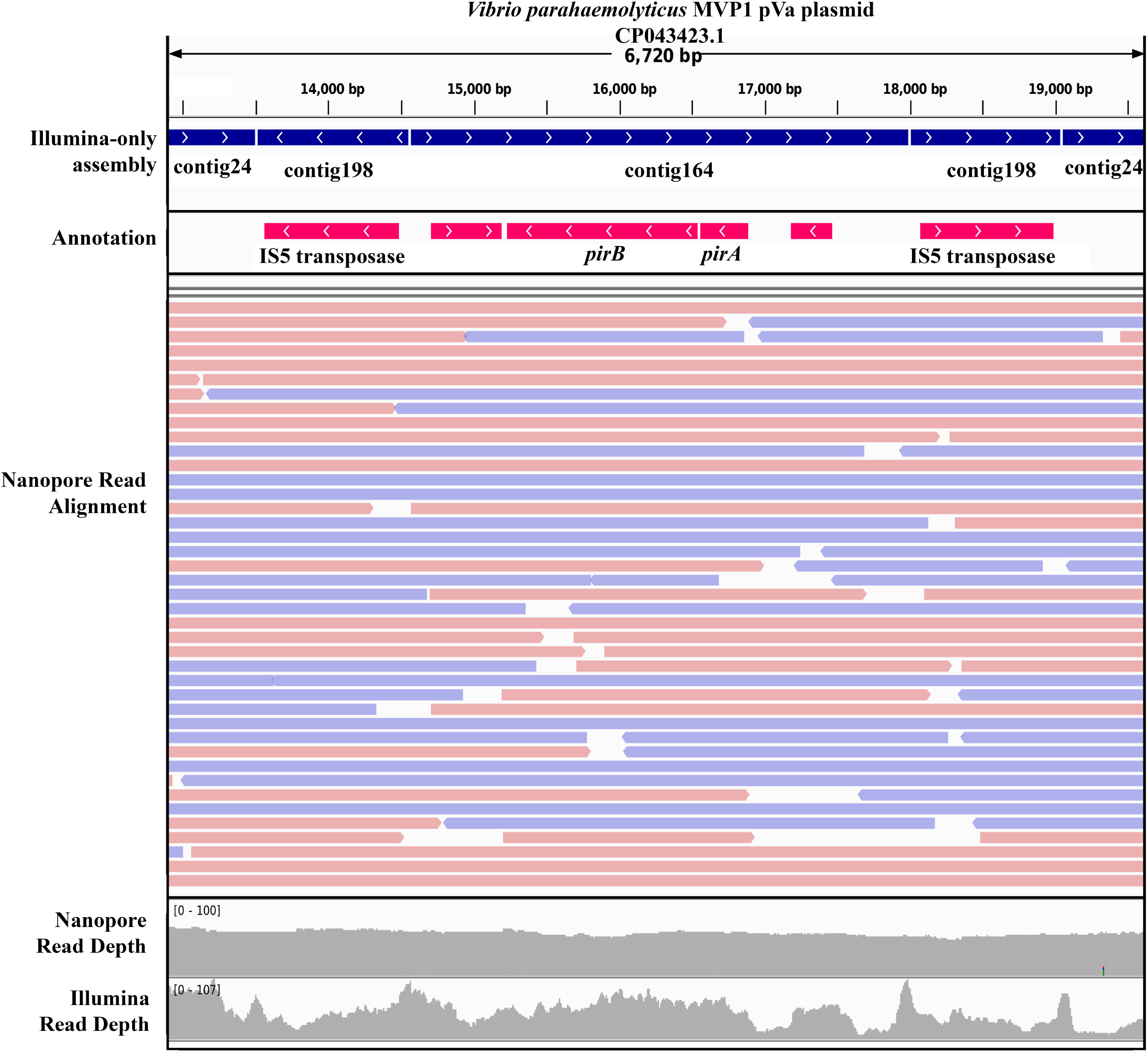
Alignment of Illumina-assembled contigs, Illumina and Nanopore reads to the annotated Vibrio *parahaemolyticus* MVP1 pVa plasmid sub-region containing the *pirAB*^Vp^ genes. Direction of arrow in the annotation indicates transcription orientation. Blue and red arrows in the Nanopore read alignment indicate forward and reverse strands, respectively.

The minor structural variations previously reported in *pirAB^Vp^*-containing region suggest that these genes are not stably maintained and are prone to transposition mediated by the flanking IS5 family transposase genes^40^. Interestingly, the localization of this IS5 family transposase gene is not specific to just the pVa plasmid. Local nucleotide similarity search of this transposase gene against the complete MVP genome revealed three additional perfect hits (100% query coverage and 100% nucleotide identity) in each of the chromosomal genome (Figure 1). The localization of these additional IS5 family transposase in both chromosome 1 and chromosome 2 further raises the possibility of the IS5 transposase-flanked *pirAB^Vp^* genes being integrated into the chromosomal region^41, 42^.

### Structural variations in the MVP1 genome

Chromosome 2 of Isolate MPV1 consists of a 160 kb genomic region that is mostly absent in five out of six pVa plasmid-harboring *V. parahaemolyticus* isolates included in this comparative genomic analysis. Functional annotation of the genes located in this region revealed two gene clusters that may further contribute to host pathogenicity in addition to the binary Pir-like toxins on the pVa plasmid. The first gene cluster located from 1,822,499 to 1,836,895 bp in chromosome 2 consists of three relatively large genes (Gene Locus Tag: BSR23_26145, BSR23_26150 and BSR23_26155) coding for another type of putative insecticidal toxins. Functional annotation using InterProScan revealed the presence of protein domains commonly associated with the toxin-complex (Tc) toxins (Figure 3). In addition, Phyre2 protein modeling also indicated their high structural homology to their respective homologous toxin components (*Extended data*: Supplemental Files 2-4). In *Xenorhabdus nematophilus*, the three toxin components formed a native toxin complex and the ingestion of this complex led to growth inhibition of two different insect larvae, *Helicoverpa zea* and *Heliothis virescens* ^43^. In addition, further tests showed that the native toxin complex is capable of binding to solubilized gut membranes of *H. zea* larvae in addition to inducing pore formation in black lipid membranes ^43^. Given the functional resemblance of such toxins to the Pir-like binary toxins, the presence of all three genes coding for the complete set of insecticidal Tc toxins in *V. parahaemolyticus* may contribute to mortality without AHPND lesions in shrimps, as previously reported ^44^.

**Figure 3.**
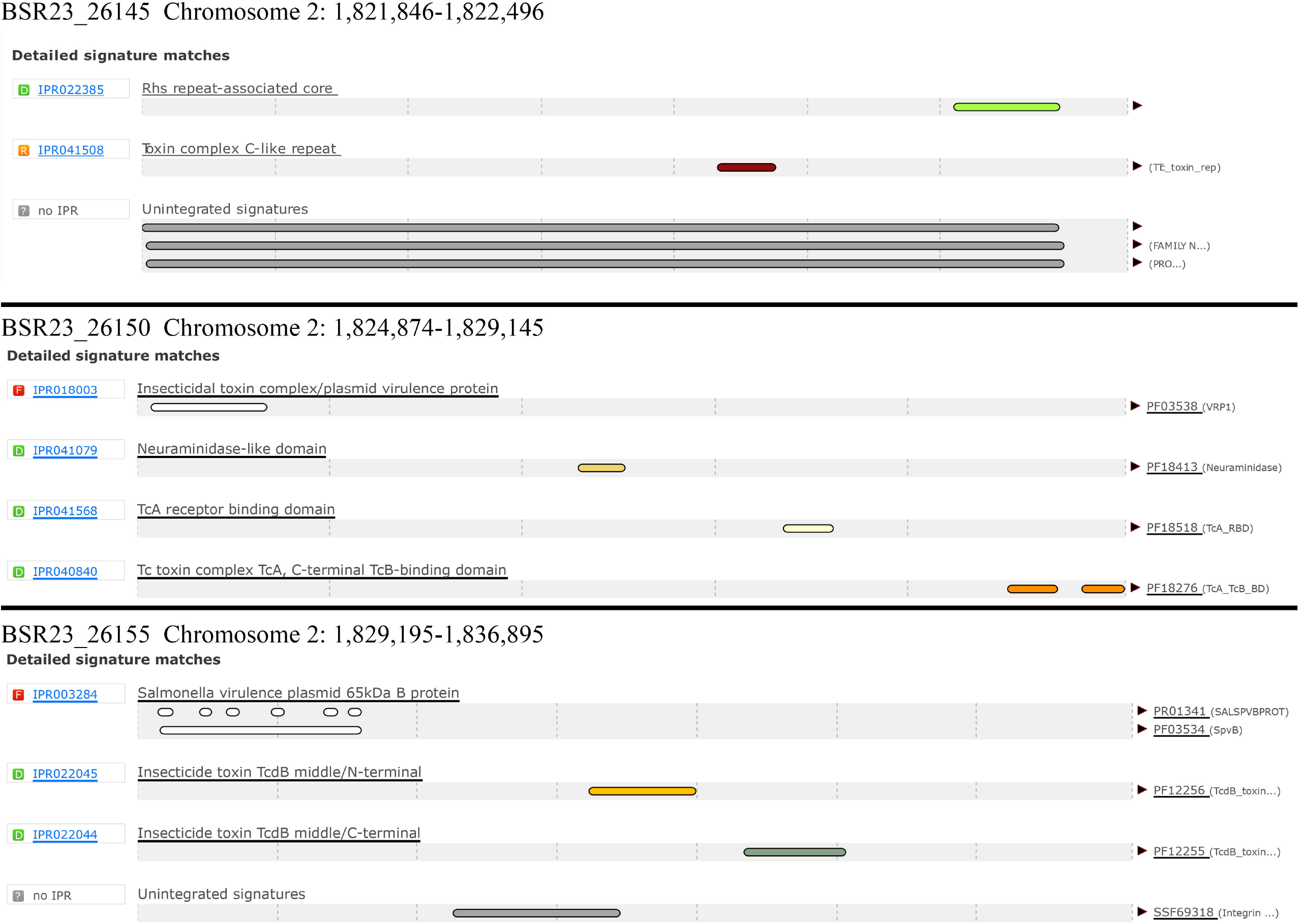
InterProScan identification of conserved protein domains in the putative toxin complex family proteins coded by three linked genes located on Chromosome 2 of *Vibrio parahaemolyticus* MVP1.

Interestingly, isolate MVP1 also harbor a *fuc* operon that is not commonly present in members of this species (*Extended data*: Supplemental Table 1). The presence of this operon in MVP1 translates into the genomic potential for the internalization of fucose for incorporation into the capsular polysaccharides or bacterial glycoproteins. While it is also possible that the *fuc* operon may contribute to improved utilization of fucose as an energy source, Williams *et al* did not observe a greater capacity to utilize fucose in an EMS-causing *V. parahaemolyticus* isolate 13-028/A3 ^45^.

### High prevalence and divergence of the LuxM/OpaM/AinS family autoinducer synthesis proteins among Vibrio spp

NCBI annotation of MVP1 proteome revealed the presence of only the LuxM/OpaM-type AHL synthase. An in-house HMMsearch for the LuxI autoinducer synthase domain (PFAM signature: PF00765) in the MVP1 proteome also did not reveal significant hits, indicating that MVP1 employs a single system for the biosynthesis of AHL signal (*Extended Data*: Supplemental File 5). Additional UniProt query search of “PF00765 AND Vibrio parahaemolyticus” on 14^th^ November 2019, showed only one positive hit across all currently available *V. parahaemolyticus* proteomes. However, further investigation of the positive hit, AAY51_09590, revealed that this belonged to a Citrobacter spp. previously misclassified as *V. parahaemolyticus* ^46^. In contrast, *V. harveyi* was shown to utilize at least two quorum-sensing systems e.g. LuxI that synthesizes 3-oxo-C6-HSL and the AinS that synthesizes C8-HSL ^47^. Despite being classified as members of the same family for sharing the similar protein domain signature, AinS exhibited a substantially lower pairwise amino acid divergence (~ 30%) in comparison to LuxM and OpaM. By rooting the LuxM/OpaM phylogenetic tree with the AinS from *Aliivibrio fischeri* as the outgroup, we observed a strongly supported monophyletic cluster consisting of the functionally validated LuxM and OpaM in addition to their homologs from other *Vibrio* spp (Figure 4A). The pairwise amino acid identity among members in this LuxM/OpaM clade averages around 50% (Figure 4B), consistent with the relatively short branch length among members of the clade.

**Figure 4.**
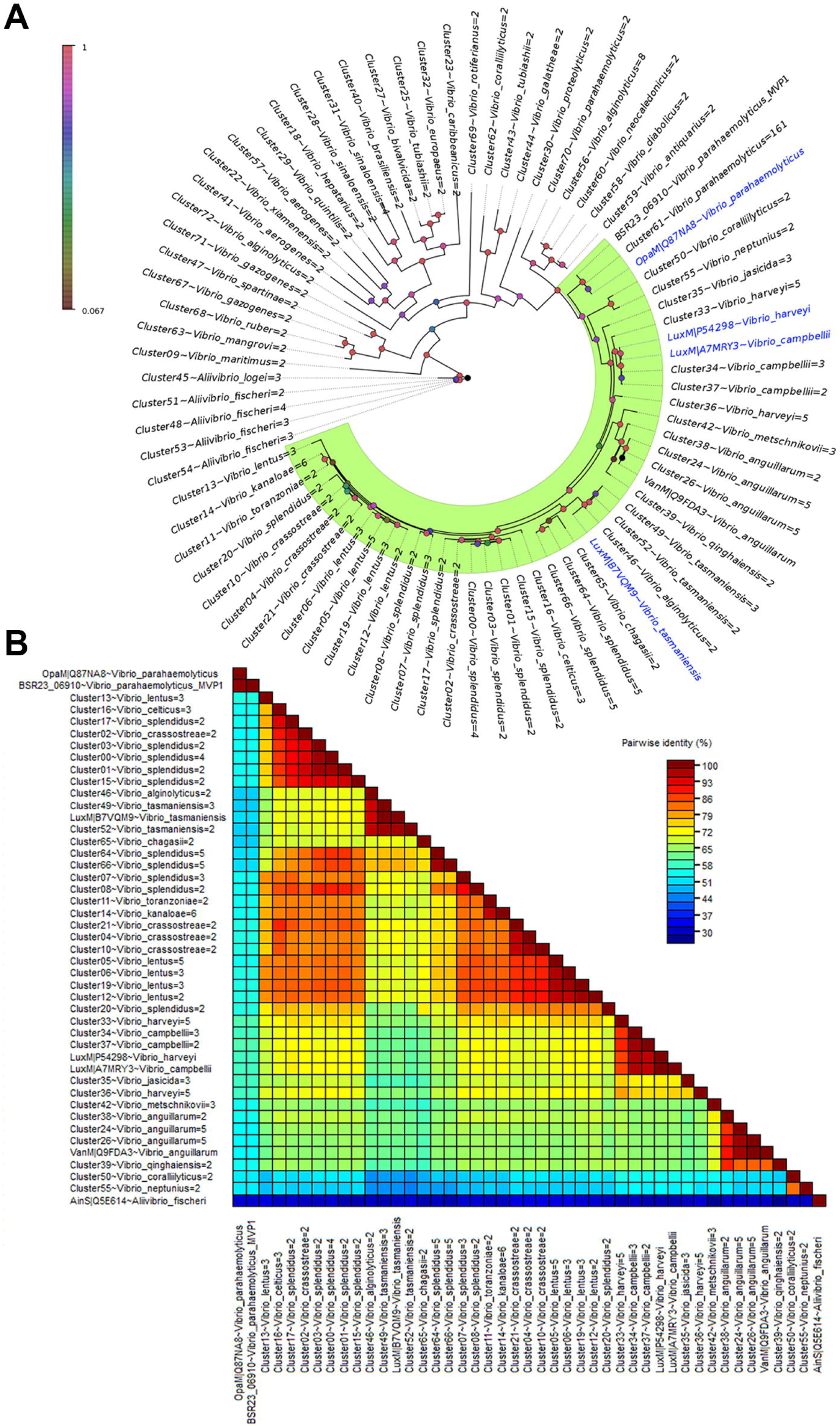
Phylogenetic analysis and sequence divergence of the AinS-like family proteins in *Vibrio* spp. (A) Maximum likelihood tree depicting the evolutionary relationships of AinS-like family protein clusters among Vibrio spp. Number after the “=” sign in each tip label indicates the number of proteins represented by the cluster. Node were colored according to the SH-like local support values and branch lengths indicate number of substitutions per site. (B) Pairwise identity matrix of the identified AinS-like family proteins in *Vibrio* spp.

In Indian white shrimps (*Fenneropenaeus indicus*), co-injection of a pathogenic *V. parahaemolyticus* strain DHAP1 and purified recombinant AHL lactonase, AiiA, reduced *Vibrio* viable counts and biofilm development in the shrimp intestine, suggesting the role of AHL-mediated quorum-sensing system in host colonization ^48^. While the presence of intact *opaM* in several sequenced *V. parahaemolyticus* isolates supports their ability to accumulate AHL and engage in quorum-sensing activity, this is best validated using a simple biosensor or LC/MS approach that can not only verify the AHL producing phenotype but also provide insight into the structural diversity of AHL signals ^49–51^. Furthermore, elucidating the role of *opaM* in mediating host colonization and pathogenesis among AHPND-causing *V. parahaemolyticus* via transposon mutagenesis will be instructive.

## Conclusions

Using approximately 40× genome coverage of Nanopore long reads, we produced the first chromosome-scale genome assembly of a Malaysian *pirAB^Vp^*-bearing *V. parahaemolyticus* isolate. Although the genome completeness of the initial Flye assembly was relatively poor, polishing of the genome assembly with Illumina reads improved its completeness to a level that is comparable to a high-quality microbial genome assembly. Structural variations identified from genomic comparisons provide new insights into the genomic features of *V. parahaemolyticus* MVP1 that may be associated with host colonization and pathogenicity.

## Data availability

### Underlying data

Vibrio parahaemolyticus strain MVP1 chromosome 1, complete sequence, Accession number CP043421: https://www.ncbi.nlm.nih.gov/nuccore/CP043421

Vibrio parahaemolyticus strain MVP1 chromosome 2, complete sequence, Accession number CP043422: https://www.ncbi.nlm.nih.gov/nuccore/CP043422

Vibrio parahaemolyticus strain MVP1 plasmid pVa, complete sequence, Accession number CP043423: https://www.ncbi.nlm.nih.gov/nuccore/CP043423

Raw Illumina reads and basecalled Nanopore Reads have also been deposited in NCBI Sequence Read Archive under the BioProject <u>PRJNA355061</u>.

### Extended data

Zenodo: Nanopore long reads enable the first complete genome assembly of a Malaysian Vibrio parahaemolyticus isolate bearing the pVa plasmid associated with acute hepatopancreatic necrosis disease,**10.5281/zenodo.3563171.** <u>Creative Commons Attribution 4.0 International</u>.

This project contains the following extended data:

Supplemental File 1: Main genome assemblies (Unpolished Flye assembly, Polished Flye assembly, Unicycler Hybrid Assembly and Unicycler Illumina-only assembly) generated in this study for comparison and their BUSCO output.

Supplemental File 2: Phyre2 protein modeling output of the putative MVP1 TcdA toxin

Supplemental File 3: Phyre2 protein modeling output of the putative MVP1 TcdB toxin

Supplemental File 4: Phyre2 protein modeling output of the putative MVP1 TccC toxin

Supplemental File 5: InterProScan output of the NCBI-predicted MVP1 proteome.

Supplemental Table 1: NCBI BlastN output using the *fuc* genes of *Vibrio parahaemolyticus* MVP1 as the query to search against the *Vibrio* reference WGS database as of 21 Oct 2019

## References

1. Haldar S, Chatterjee S, Asakura M, Vijayakumaran M, Yamasaki S. Isolation of *Vibrio parahaemolyticus* and *Vibrio cholerae* (Non-O1 and O139) from moribund shrimp (*Penaeus monodon*) and experimental challenge study against post larvae and juveniles. Annals of microbiology. 2007;57(1):55–60.

2. Lee C-T, Chen I-T, Yang Y-T, Ko T-P, Huang Y-T, Huang J-Y, et al. The opportunistic marine pathogen *Vibrio parahaemolyticus* becomes virulent by acquiring a plasmid that expresses a deadly toxin. Proceedings of the National Academy of Sciences. 2015;112(34):10798–803.

3. Yaashikaa P, Saravanan A, Kumar PS. Isolation and identification of *Vibrio cholerae* and *Vibrio parahaemolyticus* from prawn (*Penaeus monodon*) seafood: Preservation strategies. Microbial pathogenesis. 2016;99:5–13.

4. Khimmakthong U, Sukkarun P. The spread of *Vibrio parahaemolyticus* in tissues of the Pacific white shrimp *Litopenaeus vannamei* analyzed by PCR and histopathology. Microb Pathog. 2017 Dec;113:107–12. PubMed PMID: 29056496. Epub 2017/10/24. eng.

5. Soto-Rodriguez SA, Gomez-Gil B, Lozano-Olvera R, Betancourt-Lozano M, Morales-Covarrubias MS. Field and experimental evidence of *Vibrio parahaemolyticus* as the causative agent of acute hepatopancreatic necrosis disease of cultured shrimp (*Litopenaeus vannamei*) in Northwestern Mexico. Appl Environ Microbiol. 2015;81(5):1689–99.

6. Hong X, Xu D, Zhuo Y, Liu H, Lu L. Identification and pathogenicity of *Vibrio parahaemolyticus* isolates and immune responses of *Penaeus* (*Litopenaeus*) *vannamei* (Boone). Journal of fish diseases. 2016;39(9):1085–97.

7. Hong X, Lu L, Xu D. Progress in research on acute hepatopancreatic necrosis disease (AHPND). Aquaculture international. 2016;24(2):577–93.

8. Kua BC, Iar A, Siti Zahrah A, Irene J, Norazila J, Nik Haiha N, et al., editors. Current status of acute hepatopancreatic necrosis disease (AHPND) of farmed shrimp in Malaysia. Addressing Acute Hepatopancreatic Necrosis Disease (AHPND) and Other Transboundary Diseases for Improved Aquatic Animal Health in Southeast Asia: Proceedings of the ASEAN Regional Technical Consultation on EMS/AHPND and Other Transboundary Diseases for Improved Aquatic Animal Health in Southeast Asia, 22-24 February 2016, Makati City, Philippines; 2016: Aquaculture Department, Southeast Asian Fisheries Development Center.

9. Relman DA. Microbial genomics and infectious diseases. New England Journal of Medicine. 2011;365(4):347–57.

10. Yap K-P, Gan HM, Teh CSJ, Baddam R, Chai L-C, Kumar N, et al. Genome Sequence and Comparative Pathogenomics Analysis of a *Salmonella enterica* Serovar Typhi Strain Associated with a Typhoid Carrier in Malaysia. Journal of Bacteriology. 2012;194(21):5970–1.

11. Foo SM, Eng WWH, Lee YP, Gui K, Gan HM. New sequence types of *Vibrio parahaemolyticus* isolated from a Malaysian aquaculture pond, as revealed by whole-genome sequencing. Genome Announc. 2017;5(19):e00302–17.

12. Osama A, Gan HM, Teh CSJ, Yap K-P, Thong K-L. Genome sequence and comparative genomics analysis of a *Vibrio cholerae* O1 strain isolated from a cholera patient in Malaysia. Am Soc Microbiol; 2012.

13. Gan HM, Rajasekaram G, Eng WWH, Kaniappan P, Dhanoa A. Whole-Genome Sequences of Two Carbapenem-Resistant *Klebsiella quasipneumoniae* Strains Isolated from a Tertiary Hospital in Johor, Malaysia. Genome Announcements. 2017;5(32):e00768–17.

14. Gan HM, Eng WWH, Dhanoa A. Data on whole-genome sequencing of extended-spectrum beta-lactamases producing *Enterobacteriaceae* isolates from Malaysia. Data Brief. 2019;25:104257-. PubMed PMID: 31384648. eng.

15. Gan HM, Eng WWH, Dhanoa A. First genomic insights into carbapenem-resistant *Klebsiella pneumoniae* from Malaysia. Journal of Global Antimicrobial Resistance. 2019 2019/07/17/.

16. Yap K-P, Ho WS, Gan HM, Chai LC, Thong KL. Global MLST of *Salmonella* Typhi revisited in post-genomic era: genetic conservation, population structure, and comparative genomics of rare sequence types. Frontiers in microbiology. 2016;7:270.

17. Yan CZY, Austin CM, Ayub Q, Rahman S, Gan HM. Genomic characterization of *Vibrio parahaemolyticus* from Pacific white shrimp and rearing water in Malaysia reveals novel sequence types and structural variation in genomic regions containing the Photorhabdus insect-related (Pir) toxin-like genes. FEMS microbiology letters. 2019.

18. Theethakaew C, Nakamura S, Motooka D, Matsuda S, Kodama T, Chonsin K, et al. Plasmid dynamics in *Vibrio parahaemolyticus* strains related to shrimp Acute Hepatopancreatic Necrosis Syndrome (AHPNS). Infection, Genetics and Evolution. 2017;51:211–8.

19. Yang Y-T, Chen I-T, Lee C-T, Chen C-Y, Lin S-S, Hor L-I, et al. Draft genome sequences of four strains of *Vibrio parahaemolyticus*, three of which cause early mortality syndrome/acute hepatopancreatic necrosis disease in shrimp in China and Thailand. Genome Announc. 2014;2(5):e00816–14.

20. Watson M, Warr A. Errors in long-read assemblies can critically affect protein prediction. Nature Biotechnology. 2019 2019/02/01;37(2):124–6.

21. Gan HM, Lee YP, Austin CM. Nanopore long-read guided complete genome assembly of *Hydrogenophaga intermedia*, and genomic insights into 4-aminobenzenesulfonate, p-aminobenzoic acid and hydrogen metabolism in the genus *Hydrogenophaga*. Frontiers in microbiology. 2017;8:1880.

22. Austin CM, Tan MH, Harrisson KA, Lee YP, Croft LJ, Sunnucks P, et al. De novo genome assembly and annotation of Australia's largest freshwater fish, the Murray cod (*Maccullochella peelii*), from Illumina and Nanopore sequencing read. GigaScience. 2017;6(8):gix063.

23. Wick RR, Judd LM, Gorrie CL, Holt KE. Unicycler: resolving bacterial genome assemblies from short and long sequencing reads. PLoS computational biology. 2017;13(6):e1005595.

24. Kolmogorov M, Yuan J, Lin Y, Pevzner PA. Assembly of long, error-prone reads using repeat graphs. Nature biotechnology. 2019;37(5):540.

25. Walker BJ, Abeel T, Shea T, Priest M, Abouelliel A, Sakthikumar S, et al. Pilon: an integrated tool for comprehensive microbial variant detection and genome assembly improvement. PloS one. 2014;9(11):e112963.

26. Waterhouse RM, Seppey M, Simão FA, Manni M, Ioannidis P, Klioutchnikov G, et al. BUSCO applications from quality assessments to gene prediction and phylogenomics. Molecular biology and evolution. 2017;35(3):543–8.

27. Hyatt D, Chen G-L, LoCascio PF, Land ML, Larimer FW, Hauser LJ. Prodigal: prokaryotic gene recognition and translation initiation site identification. BMC bioinformatics. 2010;11(1):119.

28. Tatusova T, DiCuccio M, Badretdin A, Chetvernin V, Nawrocki EP, Zaslavsky L, et al. NCBI prokaryotic genome annotation pipeline. Nucleic acids research. 2016;44(14):6614–24.

29. Alikhan N-F, Petty NK, Zakour NLB, Beatson SA. BLAST Ring Image Generator (BRIG): simple prokaryote genome comparisons. BMC genomics. 2011;12(1):402.

30. Kelley LA, Mezulis S, Yates CM, Wass MN, Sternberg MJE. The Phyre2 web portal for protein modeling, prediction and analysis. Nat Protoc. 2015;10(6):845–58. PubMed PMID: 25950237. Epub 2015/05/07. eng.

31. Consortium U. UniProt: a hub for protein information. Nucleic acids research. 2014;43(D1):D204–D12.

32. Fu L, Niu B, Zhu Z, Wu S, Li W. CD-HIT: accelerated for clustering the next-generation sequencing data. Bioinformatics. 2012;28(23):3150–2.

33. Katoh K, Standley DM. MAFFT multiple sequence alignment software version 7: improvements in performance and usability. Molecular biology and evolution. 2013;30(4):772–80.

34. Price MN, Dehal PS, Arkin AP. FastTree 2–approximately maximum-likelihood trees for large alignments. PloS one. 2010;5(3):e9490.

35. Muhire BM, Varsani A, Martin DP. SDT: A Virus Classification Tool Based on Pairwise Sequence Alignment and Identity Calculation. PLOS ONE. 2014;9(9):e108277.

36. Gan HM, Gan HY, Ahmad NH, Aziz NA, Hudson AO, Savka MA. Whole genome sequencing and analysis reveal insights into the genetic structure, diversity and evolutionary relatedness of *luxI* and *luxR* homologs in bacteria belonging to the Sphingomonadaceae family. Frontiers in Cellular and Infection Microbiology. 2015 2015-January-08;4(188). English.

37. Krehenwinkel H, Wolf M, Lim JY, Rominger AJ, Simison WB, Gillespie RG. Estimating and mitigating amplification bias in qualitative and quantitative arthropod metabarcoding. Scientific reports. 2017;7(1):17668.

38. Gan HM, Grandjean F, Jenkins TL, Austin CM. Absence of evidence is not evidence of absence: Nanopore sequencing and complete assembly of the European lobster (*Homarus gammarus*) mitogenome uncovers the missing *nad2* and a new major gene cluster duplication. BMC genomics. 2019;20(1):335.

39. Gan HM, Linton SM, Austin CM. Two reads to rule them all: Nanopore long read-guided assembly of the iconic Christmas Island red crab, *Gecarcoidea natalis* (Pocock, 1888), mitochondrial genome and the challenges of AT-rich mitogenomes. Marine genomics. 2019;45:64–71.

40. Xiao J, Liu L, Ke Y, Li X, Liu Y, Pan Y, et al. Shrimp AHPND-causing plasmids encoding the PirAB toxins as mediated by *pirAB*-Tn903 are prevalent in various *Vibrio* species. Scientific reports. 2017;7:42177.

41. Mahillon J, Chandler M. Insertion sequences. Microbiol Mol Biol Rev. 1998;62(3):725–74.

42. Skipper KA, Andersen PR, Sharma N, Mikkelsen JG. DNA transposon-based gene vehicles - scenes from an evolutionary drive. J Biomed Sci. 2013;20(1):92-. PubMed PMID: 24320156. eng.

43. Sheets JJ, Hey TD, Fencil KJ, Burton SL, Ni W, Lang AE, et al. Insecticidal toxin complex proteins from *Xenorhabdus nematophilus* structure and pore formation. Journal of Biological Chemistry. 2011;286(26):22742–9.

44. Phiwsaiya K, Charoensapsri W, Taengphu S, Dong HT, Sangsuriya P, Nguyen GTT, et al. A Natural V*ibrio parahaemolyticus* DeltapirA (Vp) pirB (Vp+) Mutant Kills Shrimp but Produces neither Pir (Vp) Toxins nor Acute Hepatopancreatic Necrosis Disease Lesions. Applied and environmental microbiology. 2017 Aug 15;83(16). PubMed PMID: 28576761. Pubmed Central PMCID: PMC5541212. Epub 2017/06/04. eng.

45. Williams SL, Jensen RV, Kuhn DD, Stevens AM. Analyzing the metabolic capabilities of a *Vibrio parahaemolyticus* strain that causes early mortality syndrome in shrimp. Aquaculture. 2017;476:44–8.

46. Allnutt T, Yan CZY, Crowley TM, Gan HM. Commentary: genome sequence of *Vibrio parahaemolyticus* VP152 strain isolated from *Penaeus indicus* in malaysia. Frontiers in microbiology. 2018;9:865.

47. Gilson L, Kuo A, Dunlap PV. AinS and a new family of autoinducer synthesis proteins. J Bacteriol. 1995 Dec;177(23):6946–51. PubMed PMID: 7592489. Pubmed Central PMCID: PMC177564. Epub 1995/12/01. eng.

48. Vinoj G, Vaseeharan B, Thomas S, Spiers AJ, Shanthi S. Quorum-Quenching Activity of the AHL-Lactonase from *Bacillus licheniformis* DAHB1 Inhibits *Vibrio* Biofilm Formation *In Vitro* and Reduces Shrimp Intestinal Colonisation and Mortality. Marine Biotechnology. 2014 December 01;16(6):707–15.

49. Gan HM, Dailey LK, Halliday N, Williams P, Hudson AO, Savka MA. Genome sequencing-assisted identification and the first functional validation of N-acyl-homoserine-lactone synthases from the Sphingomonadaceae family. PeerJ. 2016 2016/08/30;4:e2332.

50. Gan HM, Buckley L, Szegedi E, Hudson AO, Savka MA. Identification of an *rsh* Gene from a *Novosphingobium* sp. Necessary for Quorum-Sensing Signal Accumulation. Journal of Bacteriology. 2009;191(8):2551–60.

51. Lowe N, Gan HM, Chakravartty V, Scott R, Szegedi E, Burr TJ, et al. Quorum-sensing signal production by *Agrobacterium vitis* strains and their tumor-inducing and tartrate-catabolic plasmids. FEMS Microbiology Letters. 2009;296(1):102–9.

