## Supplementary material for "Nanopore long reads enable the first complete genome assembly of a Malaysian *Vibrio parahaemolyticus* isolate bearing the pVa plasmid associated with acute hepatopancreatic necrosis disease": Supplmental File 1 (from another version of this preprint: version 1, 2019-12-02)

Phyre2

| | |
|---|---|
| Email | |
| Description | Undefined |
| Date | Tue Oct 22 03:04:19 BST 2019 |
| Unique Job ID | 6f83532eb032a780 |

Detailed template information

| # | Template | Alignment Coverage | 3D Model | Confidence | % i.d. | Template Information |
|---|---|---|---|---|---|---|
| 1 | c4o9xA_ | Alignment | 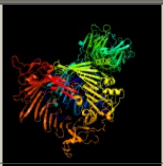 | 100.0 | 50 | **PDB header:**toxin<br>**Chain:** A: **PDB Molecule:**tcdb2, tccc3;<br>**PDBTitle:** crystal structure of tcdb2-tccc3 |
| 2 | c4iglA_ | Alignment | 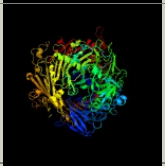 | 100.0 | 48 | **PDB header:**toxin<br>**Chain:** A: **PDB Molecule:**yenb;<br>**PDBTitle:** structure of the rhs-repeat containing bc component of the secreted2 abc toxin complex from yersinia entomophaga |
| 3 | c6p0xA_ | Alignment | 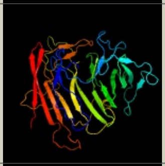 | 100.0 | 45 | **PDB header:**unknown function<br>**Chain:** A: **PDB Molecule:**salmonella plasmid virulence: spvb;<br>**PDBTitle:** structure of the n-terminal domain of effector protein spvb from2 salmonella typhimurium strain lt2 |
| 4 | c6fb3A_ | Alignment | 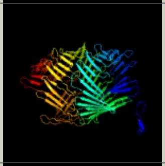 | 99.9 | 12 | **PDB header:**cell adhesion<br>**Chain:** A: **PDB Molecule:**teneurin-2;<br>**PDBTitle:** teneurin 2 partial extracellular domain |
| 5 | c4tqjA_ | Alignment | 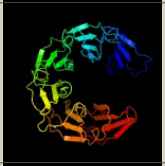 | 99.9 | 17 | **PDB header:**sugar binding protein<br>**Chain:** A: **PDB Molecule:**lectin 2;<br>**PDBTitle:** structural basis of specific recognition of non-reducing terminal n-2 acetylglucosamine by an agrocybe aegerita lection |
| 6 | c2c4dA_ | Alignment | 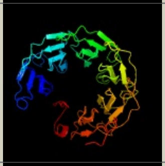 | 99.9 | 15 | **PDB header:**lectin<br>**Chain:** A: **PDB Molecule:**psathyrella velutina lectin pvl;<br>**PDBTitle:** 2.6a crystal structure of psathyrella velutina lectin in2 complex with n-acetylglucosamine |
| 7 | c6fayA_ | Alignment | 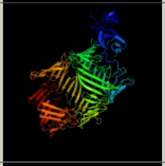 | 99.9 | 11 | **PDB header:**cell adhesion<br>**Chain:** A: **PDB Molecule:**odz3 protein;<br>**PDBTitle:** teneurin3 monomer |
| 8 | c4iglB_ | Alignment | 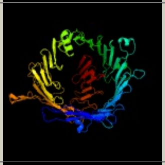 | 99.9 | 11 | **PDB header:**toxin<br>**Chain:** B: **PDB Molecule:**yenc2;<br>**PDBTitle:** structure of the rhs-repeat containing bc component of the secreted2 abc toxin complex from yersinia entomophaga |
| 9 | c5kisB_ | Alignment | 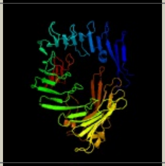 | 99.9 | 13 | **PDB header:**toxin<br>**Chain:** B: **PDB Molecule:**rhs2;<br>**PDBTitle:** yenb/rhs2 complex |
| 10 | c6cmxA_ | Alignment | 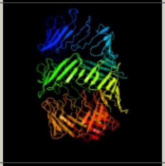 | 99.9 | 11 | **PDB header:**membrane protein<br>**Chain:** A: **PDB Molecule:**teneurin-2;<br>**PDBTitle:** human teneurin 2 extra-cellular region |
| 11 | c4ghbA_ | Alignment | 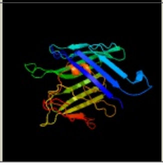 | 99.6 | 10 | **PDB header:**structural genomics, unknown function<br>**Chain:** A: **PDB Molecule:**hypothetical protein;<br>**PDBTitle:** crystal structure of a porin-like protein (bacuni_01323) from2 bacteroides uniformis atcc 8492 at 2.32 a resolution |

| | | | | | | |
|---|---|---|---|---|---|---|
| 12 | c4a7kA_ | Alignment | 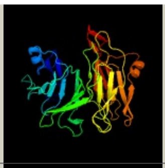 | 99.5 | 15 | **PDB header:**lyase<br>**Chain:** A: **PDB Molecule:**aldos-2-ulose dehydratase;<br>**PDBTitle:** bifunctional aldos-2-ulose dehydratase |
| 13 | c6mltA_ | Alignment | 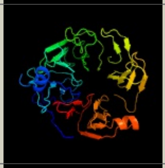 | 99.4 | 24 | **PDB header:**sugar binding protein<br>**Chain:** A: **PDB Molecule:**hemolysin-related protein;<br>**PDBTitle:** crystal structure of the v. cholerae biofilm matrix protein bap1 |
| 14 | c4g1eA_ | Alignment | 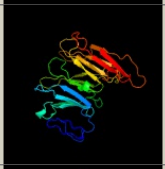 | 99.1 | 17 | **PDB header:**protein binding<br>**Chain:** A: **PDB Molecule:**integrin alpha-v;<br>**PDBTitle:** crystal structure of integrin alpha v beta 3 with coil-coiled tag. |
| 15 | c2zuxA_ | Alignment | 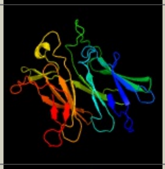 | 99.0 | 15 | **PDB header:**lyase<br>**Chain:** A: **PDB Molecule:**yesw protein;<br>**PDBTitle:** crystal structure of rhamnogalacturonan lyase yesw2 complexed with rhamnose |
| 16 | c2zuyA_ | Alignment | 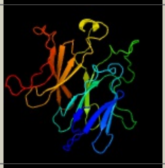 | 99.0 | 14 | **PDB header:**lyase<br>**Chain:** A: **PDB Molecule:**yesx protein;<br>**PDBTitle:** crystal structure of exotype rhamnogalacturonan lyase yesx |
| 17 | c3vi4C_ | Alignment | 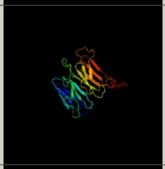 | 98.9 | 17 | **PDB header:**cell adhesion/immune system<br>**Chain:** C: **PDB Molecule:**integrin alpha-5;<br>**PDBTitle:** crystal structure of alpha5beta1 integrin headpiece in complex with2 rgd peptide |
| 18 | c1l5gA_ | Alignment | 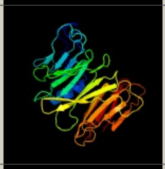 | 98.9 | 13 | **PDB header:**cell adhesion<br>**Chain:** A: **PDB Molecule:**integrin alpha v;<br>**PDBTitle:** crystal structure of the extracellular segment of integrin avb3 in2 complex with an arg-gly-asp ligand |
| 19 | c3k6sC_ | Alignment | 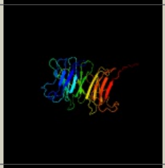 | 98.9 | 14 | **PDB header:**cell adhesion<br>**Chain:** C: **PDB Molecule:**integrin alpha-x;<br>**PDBTitle:** structure of integrin alphaxbeta2 ectodomain |
| 20 | c5e6rA_ | Alignment | 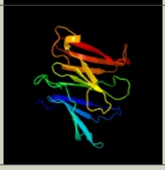 | 98.8 | 17 | **PDB header:**cell adhesion<br>**Chain:** A: **PDB Molecule:**integrin alpha-l;<br>**PDBTitle:** structures of leukocyte integrin alb2: the ai domain, the headpiece,2 and the pocket for the internal ligand |
| 21 | c3v4vA_ | Alignment | not modelled | 98.8 | 11 | **PDB header:**cell adhesion<br>**Chain:** A: **PDB Molecule:**integrin alpha-4;<br>**PDBTitle:** crystal structure of a4b7 headpiece complexed with fab act-1 and2 ro0505376 |
| 22 | c3fcsA_ | Alignment | not modelled | 98.7 | 17 | **PDB header:**cell adhesion/blood clotting<br>**Chain:** A: **PDB Molecule:**integrin, alpha 2b;<br>**PDBTitle:** structure of complete ectodomain of integrin aiibb3 |
| 23 | c3k71G_ | Alignment | not modelled | 98.5 | 15 | **PDB header:**cell adhesion<br>**Chain:** G: **PDB Molecule:**integrin alpha-x;<br>**PDBTitle:** structure of integrin alphax beta2 ectodomain |
| 24 | d1tyea_ | Alignment | not modelled | 98.5 | 18 | **Fold:**7-bladed beta-propeller<br>**Superfamily:**Integrin alpha N-terminal domain<br>**Family:**Integrin alpha N-terminal domain |
| 25 | d1jv2a4 | Alignment | not modelled | 98.4 | 14 | **Fold:**7-bladed beta-propeller<br>**Superfamily:**Integrin alpha N-terminal domain<br>**Family:**Integrin alpha N-terminal domain |
| 26 | c5mxhA_ | Alignment | not modelled | 97.8 | 11 | **PDB header:**sugar binding protein<br>**Chain:** A: **PDB Molecule:**photorhabdus asymbiotica lectin phl;<br>**PDBTitle:** photorhabdus asymbiotica lectin (phl) in complex with d-galactose |
| 27 | c5c9pA_ | Alignment | not modelled | 96.9 | 8 | **PDB header:**sugar binding protein<br>**Chain:** A: **PDB Molecule:**pll lectin;<br>**PDBTitle:** crystal structure of recombinant pll lectin complexed with l-fucose2 from photorhabdus luminescens at 1.75 a resolution |
| 28 | c4v0nH_ | Alignment | not modelled | 67.3 | 10 | **PDB header:**hydrolase/structural protein<br>**Chain:** H: **PDB Molecule:**bardet-biedl syndrome 1 protein;<br>**PDBTitle:** crystal structure of bbs1n in complex with arl6dn, soaked with mercury |
| | | | | | | **PDB header:**ligase |

| | | | | | | |
|---|---|---|---|---|---|---|
| 29 | c4rohA_ | Alignment | not modelled | 63.2 | 13 | **Chain:** A: **PDB Molecule:**e3 ubiquitin-protein ligase itchy homolog; **PDBTitle:** crystal structure of tandem ww domains of itch in complex with txnip2 peptide |
| 30 | c2b59B_ | Alignment | not modelled | 59.7 | 7 | **PDB header:**hydrolase/structural protein **Chain:** B: **PDB Molecule:**cellulosomal scaffolding protein a; **PDBTitle:** the type ii cohesin dockerin complex |
| 31 | c4qtqA_ | Alignment | not modelled | 52.7 | 28 | **PDB header:**hydrolase inhibitor **Chain:** A: **PDB Molecule:**xac2610 protein; **PDBTitle:** structure of a xanthomonas type iv secretion system related protein |
| 32 | c4qi3A_ | Alignment | not modelled | 40.3 | 17 | **PDB header:**oxidoreductase **Chain:** A: **PDB Molecule:**cellobiose dehydrogenase; **PDBTitle:** cytochrome domain of myriococcum thermophilum cellobiose2 dehydrogenase, mtcyt |
| 33 | c2l5fA_ | Alignment | not modelled | 37.6 | 17 | **PDB header:**protein binding **Chain:** A: **PDB Molecule:**pre-mrna-processing factor 40 homolog a; **PDBTitle:** solution structure of the tandem ww domains from hypa/fbp11 |
| 34 | c2kxqA_ | Alignment | not modelled | 37.4 | 18 | **PDB header:**protein binding **Chain:** A: **PDB Molecule:**e3 ubiquitin-protein ligase smurf2; **PDBTitle:** solution structure of smurf2 ww2 and ww3 bound to smad7 py motif2 containing peptide |
| 35 | c3qq2C_ | Alignment | not modelled | 32.8 | 11 | **PDB header:**membrane protein/protein transport **Chain:** C: **PDB Molecule:**brka autotransporter; **PDBTitle:** crystal structure of the beta domain of the bordetella autotransporter2 brka |
| 36 | d2dk1a1 | Alignment | not modelled | 32.8 | 10 | **Fold:**WW domain-like **Superfamily:**WW domain **Family:**WW domain |
| 37 | c1tk7A_ | Alignment | not modelled | 30.4 | 19 | **PDB header:**signaling protein **Chain:** A: **PDB Molecule:**cg4244-pb; **PDBTitle:** nmr structure of ww domains (ww3-4) from suppressor of2 deltex |
| 38 | c3kcpA_ | Alignment | not modelled | 28.9 | 13 | **PDB header:**structural protein **Chain:** A: **PDB Molecule:**cellulosomal-scaffolding protein a; **PDBTitle:** crystal structure of interacting clostridium thermocellum2 multimodular components |
| 39 | c4zgfA_ | Alignment | not modelled | 26.8 | 19 | **PDB header:**unknown function **Chain:** A: **PDB Molecule:**uncharacterized protein; **PDBTitle:** crystal structure of a duf4847 family protein (bvu_2626) from2 bacteroides vulgatus atcc 8482 at 1.00 a resolution |
| 40 | d1lgta2 | Alignment | not modelled | 26.4 | 6 | **Fold:**Glyoxalase/Bleomycin resistance protein/Dihydroxybiphenyl dioxygenase **Superfamily:**Glyoxalase/Bleomycin resistance protein/Dihydroxybiphenyl dioxygenase **Family:**Extradiol dioxygenases |
| 41 | c1zr7A_ | Alignment | not modelled | 21.1 | 11 | **PDB header:**signaling protein **Chain:** A: **PDB Molecule:**huntingtin-interacting protein hypa/fbp11; **PDBTitle:** solution structure of the first ww domain of fbp11 |
| 42 | c4huzA_ | Alignment | not modelled | 20.6 | 5 | **PDB header:**oxidoreductase **Chain:** A: **PDB Molecule:**2,6-dichloro-p-hydroquinone 1,2-dioxygenase; **PDBTitle:** 2,6-dichloro-p-hydroquinone 1,2-dioxygenase |
| 43 | d1o6wa1 | Alignment | not modelled | 19.9 | 4 | **Fold:**WW domain-like **Superfamily:**WW domain **Family:**WW domain |
| 44 | c6jjzA_ | Alignment | not modelled | 19.9 | 12 | **PDB header:**signaling protein **Chain:** A: **PDB Molecule:**membrane-associated guanylate kinase, ww and pdz domain- **PDBTitle:** crystal structure of magi2 and dendrin complex |
| 45 | c3rheA_ | Alignment | not modelled | 19.7 | 4 | **PDB header:**oxidoreductase **Chain:** A: **PDB Molecule:**nad-dependent benzaldehyde dehydrogenase; **PDBTitle:** the crystal structure of nad-dependent benzaldehyde dehydrogenase from2 legionella pneumophila |
| 46 | c2dk1A_ | Alignment | not modelled | 18.7 | 14 | **PDB header:**gene regulation **Chain:** A: **PDB Molecule:**ww domain-binding protein 4; **PDBTitle:** solution structure of ww domain in ww domain binding2 protein 4 (wbp-4) |
| 47 | c3fcdB_ | Alignment | not modelled | 18.6 | 15 | **PDB header:**lyase **Chain:** B: **PDB Molecule:**lyase; **PDBTitle:** crystal structure of a putative glyoxalase from an2 environmental bacteria |
| 48 | d1kw3b1 | Alignment | not modelled | 18.4 | 12 | **Fold:**Glyoxalase/Bleomycin resistance protein/Dihydroxybiphenyl dioxygenase **Superfamily:**Glyoxalase/Bleomycin resistance protein/Dihydroxybiphenyl dioxygenase **Family:**Extradiol dioxygenases |
| 49 | d1jifa_ | Alignment | not modelled | 17.9 | 8 | **Fold:**Glyoxalase/Bleomycin resistance protein/Dihydroxybiphenyl dioxygenase **Superfamily:**Glyoxalase/Bleomycin resistance protein/Dihydroxybiphenyl dioxygenase **Family:**Antibiotic resistance proteins |
| 50 | c5ii6A_ | Alignment | not modelled | 17.7 | 38 | **PDB header:**cell adhesion **Chain:** A: **PDB Molecule:**zona pellucida sperm-binding protein 2; **PDBTitle:** crystal structure of the zp-n1 domain of mouse sperm receptor zp2 at2 0.95 a resolution |
| 51 | c3e5dA_ | Alignment | not modelled | 17.4 | 11 | **PDB header:**lyase **Chain:** A: **PDB Molecule:**putative glyoxalase i; **PDBTitle:** crystal structure of a putative glyoxalase i (lmof2365_0426) from2 listeria monocytogenes str. 4b f2365 at 2.70 a resolution |
| 52 | c2mweA_ | Alignment | not modelled | 17.3 | 13 | **PDB header:**transcription **Chain:** A: **PDB Molecule:**transcription elongation regulator 1; **PDBTitle:** nmr structure of fbp28 ww2 mutant y438r, l453a dndc |
| 53 | c2mwfA_ | Alignment | not modelled | 15.8 | 11 | **PDB header:**transcription **Chain:** A: **PDB Molecule:**transcription elongation regulator 1; **PDBTitle:** nmr structure of fbp28 ww2 mutant y438r dn |
| | | | | | | **Fold:**Glyoxalase/Bleomycin resistance protein/Dihydroxybiphenyl |

| | | | | | | |
|---|---|---|---|---|---|---|
| 54 | d1f1ua2 | Alignment | not modelled | 15.6 | 13 | dioxygenase<br>**Superfamily:**Glyoxalase/Bleomycin resistance protein/Dihydroxybiphenyl dioxygenase<br>**Family:**Extradiol dioxygenases |
| 55 | c4z04A_ | Alignment | not modelled | 15.6 | 14 | **PDB header:**lyase<br>**Chain:** A: **PDB Molecule:**glyoxalase/bleomycin resistance /dioxygenase superfamily<br>**PDBTitle:** crystal structure of a probable lactoylglutathione lyase from brucella2 melitensis in complex with glutathione |
| 56 | c3hx6A_ | Alignment | not modelled | 15.6 | 25 | **PDB header:**cell adhesion<br>**Chain:** A: **PDB Molecule:**type 4 fimbrial biogenesis protein pily1;<br>**PDBTitle:** crystal structure of pseudomonas aeruginosa pily1 c-terminal2 domain |
| 57 | d1lgta1 | Alignment | not modelled | 15.4 | 14 | **Fold:**Glyoxalase/Bleomycin resistance protein/Dihydroxybiphenyl dioxygenase<br>**Superfamily:**Glyoxalase/Bleomycin resistance protein/Dihydroxybiphenyl dioxygenase<br>**Family:**Extradiol dioxygenases |
| 58 | c6j69A_ | Alignment | not modelled | 15.1 | 11 | **PDB header:**cell cycle<br>**Chain:** A: **PDB Molecule:**protein kibra;<br>**PDBTitle:** structure of kibra and dendrin complex |
| 59 | c3r6aB_ | Alignment | not modelled | 14.9 | 17 | **PDB header:**isomerase, lyase<br>**Chain:** B: **PDB Molecule:**uncharacterized protein;<br>**PDBTitle:** crystal structure of an uncharacterized protein (hypothetical protein2 mm_3218) from methanosarcina mazei. |
| 60 | d1f1xa2 | Alignment | not modelled | 14.9 | 7 | **Fold:**Glyoxalase/Bleomycin resistance protein/Dihydroxybiphenyl dioxygenase<br>**Superfamily:**Glyoxalase/Bleomycin resistance protein/Dihydroxybiphenyl dioxygenase<br>**Family:**Extradiol dioxygenases |
| 61 | c2mwdA_ | Alignment | not modelled | 14.7 | 13 | **PDB header:**transcription<br>**Chain:** A: **PDB Molecule:**transcription elongation regulator 1;<br>**PDBTitle:** nmr structure of fbp28 ww2 mutant y438r dndc |
| 62 | c2dk7A_ | Alignment | not modelled | 14.5 | 18 | **PDB header:**transcription<br>**Chain:** A: **PDB Molecule:**transcription elongation regulator 1;<br>**PDBTitle:** solution structure of ww domain in transcription elongation2 regulator 1 |
| 63 | c6jk0A_ | Alignment | not modelled | 14.4 | 18 | **PDB header:**signaling protein<br>**Chain:** A: **PDB Molecule:**transcriptional coactivator yap1,dendrin;<br>**PDBTitle:** crystal structure of yap1 and dendrin complex |
| 64 | d1kw3b2 | Alignment | not modelled | 14.3 | 0 | **Fold:**Glyoxalase/Bleomycin resistance protein/Dihydroxybiphenyl dioxygenase<br>**Superfamily:**Glyoxalase/Bleomycin resistance protein/Dihydroxybiphenyl dioxygenase<br>**Family:**Extradiol dioxygenases |
| 65 | d1w0pa1 | Alignment | not modelled | 14.3 | 20 | **Fold:**Concanavalin A-like lectins/glucanases<br>**Superfamily:**Concanavalin A-like lectins/glucanases<br>**Family:**Vibrio cholerae sialidase, N-terminal and insertion domains |
| 66 | d1zr7a1 | Alignment | not modelled | 14.2 | 13 | **Fold:**WW domain-like<br>**Superfamily:**WW domain<br>**Family:**WW domain |
| 67 | d1r9ca_ | Alignment | not modelled | 14.1 | 0 | **Fold:**Glyoxalase/Bleomycin resistance protein/Dihydroxybiphenyl dioxygenase<br>**Superfamily:**Glyoxalase/Bleomycin resistance protein/Dihydroxybiphenyl dioxygenase<br>**Family:**Antibiotic resistance proteins |
| 68 | d1f1xa1 | Alignment | not modelled | 13.9 | 6 | **Fold:**Glyoxalase/Bleomycin resistance protein/Dihydroxybiphenyl dioxygenase<br>**Superfamily:**Glyoxalase/Bleomycin resistance protein/Dihydroxybiphenyl dioxygenase<br>**Family:**Extradiol dioxygenases |
| 69 | c5cb9A_ | Alignment | not modelled | 13.9 | 15 | **PDB header:**lyase<br>**Chain:** A: **PDB Molecule:**glyoxalase/bleomycin resistance protein/dioxygenase;<br>**PDBTitle:** crystal structure of c-as lyase with mercaptoethonal |
| 70 | d1o6wa2 | Alignment | not modelled | 13.7 | 18 | **Fold:**WW domain-like<br>**Superfamily:**WW domain<br>**Family:**WW domain |
| 71 | c2ysiA_ | Alignment | not modelled | 13.6 | 25 | **PDB header:**protein binding<br>**Chain:** A: **PDB Molecule:**transcription elongation regulator 1;<br>**PDBTitle:** solution structure of the first ww domain from the mouse2 transcription elongation regulator 1, transcription factor3 ca150 |
| 72 | d1xrka_ | Alignment | not modelled | 13.5 | 8 | **Fold:**Glyoxalase/Bleomycin resistance protein/Dihydroxybiphenyl dioxygenase<br>**Superfamily:**Glyoxalase/Bleomycin resistance protein/Dihydroxybiphenyl dioxygenase<br>**Family:**Antibiotic resistance proteins |
| 73 | d1ywia1 | Alignment | not modelled | 13.4 | 13 | **Fold:**WW domain-like<br>**Superfamily:**WW domain<br>**Family:**WW domain |
| 74 | c1ywjA_ | Alignment | not modelled | 13.4 | 13 | **PDB header:**structural protein<br>**Chain:** A: **PDB Molecule:**formin-binding protein 3;<br>**PDBTitle:** structure of the fbp11ww1 domain |
| 75 | d1f1ua1 | Alignment | not modelled | 13.4 | 18 | **Fold:**Glyoxalase/Bleomycin resistance protein/Dihydroxybiphenyl dioxygenase<br>**Superfamily:**Glyoxalase/Bleomycin resistance protein/Dihydroxybiphenyl dioxygenase<br>**Family:**Extradiol dioxygenases |
| 76 | c5vb0E_ | Alignment | not modelled | 13.1 | 6 | **PDB header:**transferase<br>**Chain:** E: **PDB Molecule:**fosfomycin resistance protein fosa3;<br>**PDBTitle:** crystal structure of fosfomycin resistance protein fosa3 |
| 77 | d2i7ra1 | Alignment | not modelled | 12.9 | 6 | **Fold:**Glyoxalase/Bleomycin resistance protein/Dihydroxybiphenyl dioxygenase<br>**Superfamily:**Glyoxalase/Bleomycin resistance protein/Dihydroxybiphenyl dioxygenase<br>**Family:**Antibiotic resistance proteins |

| | | | | | | |
|---|---|---|---|---|---|---|
| 78 | c2jxwA_ | Alignment | not modelled | 12.8 | 11 | **PDB header:**formin binding protein<br>**Chain:** A: **PDB Molecule:**ww domain-binding protein 4;<br>**PDBTitle:** solution structure of the tandem ww domains of fbp21 |
| 79 | c6on3C_ | Alignment | not modelled | 12.7 | 26 | **PDB header:**oxidoreductase<br>**Chain:** C: **PDB Molecule:**l-dopa extradiol dioxygenase;<br>**PDBTitle:** a substrate bound structure of l-dopa dioxygenase from streptomyces2 sclerotialus |
| 80 | d1mpya1 | Alignment | not modelled | 12.6 | 12 | **Fold:**Glyoxalase/Bleomycin resistance protein/Dihydroxybiphenyl dioxygenase<br>**Superfamily:**Glyoxalase/Bleomycin resistance protein/Dihydroxybiphenyl dioxygenase<br>**Family:**Extradiol dioxygenases |
| 81 | d1q0oa2 | Alignment | not modelled | 12.6 | 8 | **Fold:**Glyoxalase/Bleomycin resistance protein/Dihydroxybiphenyl dioxygenase<br>**Superfamily:**Glyoxalase/Bleomycin resistance protein/Dihydroxybiphenyl dioxygenase<br>**Family:**Extradiol dioxygenases |
| 82 | c6iptB_ | Alignment | not modelled | 12.3 | 8 | **PDB header:**transferase<br>**Chain:** B: **PDB Molecule:**all3014 protein;<br>**PDBTitle:** crystal structure of a fosfomycin and bleomycin resistant protein from2 anabaena/nostoc cyanobacterium at 1.70 a resolution |
| 83 | c4nb2B_ | Alignment | not modelled | 12.3 | 7 | **PDB header:**transferase<br>**Chain:** B: **PDB Molecule:**metallothiol transferase fosb;<br>**PDBTitle:** crystal structure of fosb from staphylococcus aureus at 1.89 angstrom2 resolution - apo structure |
| 84 | d12asa_ | Alignment | not modelled | 12.0 | 11 | **Fold:**Class II aaRS and biotin synthetases<br>**Superfamily:**Class II aaRS and biotin synthetases<br>**Family:**Class II aminoacyl-tRNA synthetase (aaRS)-like, catalytic domain |
| 85 | c3bt3B_ | Alignment | not modelled | 11.9 | 15 | **PDB header:**structural genomics, unknown function<br>**Chain:** B: **PDB Molecule:**glyoxalase-related enzyme, arac type;<br>**PDBTitle:** crystal structure of a glyoxalase-related enzyme from clostridium2 phytofermentans |
| 86 | d2rm0w1 | Alignment | not modelled | 11.8 | 11 | **Fold:**WW domain-like<br>**Superfamily:**WW domain<br>**Family:**WW domain |
| 87 | c2qqzB_ | Alignment | not modelled | 11.8 | 15 | **PDB header:**structural genomics, unknown function<br>**Chain:** B: **PDB Molecule:**glyoxalase family protein, putative;<br>**PDBTitle:** crystal structure of putative glyoxalase family protein from bacillus2 anthracis |
| 88 | c4qi7A_ | Alignment | not modelled | 11.7 | 17 | **PDB header:**oxidoreductase<br>**Chain:** A: **PDB Molecule:**cellobiose dehydrogenase;<br>**PDBTitle:** cellobiose dehydrogenase from neurospora crassa, nccdh |
| 89 | c3aehB_ | Alignment | not modelled | 11.7 | 13 | **PDB header:**hydrolase<br>**Chain:** B: **PDB Molecule:**hemoglobin-binding protease hbp autotransporter;<br>**PDBTitle:** integral membrane domain of autotransporter hbp |
| 90 | d1pl3a_ | Alignment | not modelled | 11.5 | 12 | **Fold:**Immunoglobulin-like beta-sandwich<br>**Superfamily:**CBD9-like<br>**Family:**Cytochrome domain of cellobiose dehydrogenase |
| 91 | c2wl9A_ | Alignment | not modelled | 11.4 | 6 | **PDB header:**oxidoreductase<br>**Chain:** A: **PDB Molecule:**catechol 2,3-dioxygenase;<br>**PDBTitle:** crystal structure of catechol 2,3-dioxygenase |
| 92 | d1rk8c_ | Alignment | not modelled | 11.4 | 23 | **Fold:**WW domain-like<br>**Superfamily:**Pym (Within the bgcn gene intron protein, WIBG), N-terminal domain<br>**Family:**Pym (Within the bgcn gene intron protein, WIBG), N-terminal domain |
| 93 | c1rk8C_ | Alignment | not modelled | 11.4 | 23 | **PDB header:**translation<br>**Chain:** C: **PDB Molecule:**within the bgcn gene intron protein;<br>**PDBTitle:** structure of the cytosolic protein pym bound to the mago-2 y14 core of the exon junction complex |
| 94 | c3zw5A_ | Alignment | not modelled | 11.2 | 7 | **PDB header:**lyase<br>**Chain:** A: **PDB Molecule:**glyoxalase domain-containing protein 5;<br>**PDBTitle:** crystal structure of the human glyoxalase domain-containing protein 5 |
| 95 | d1vhoa1 | Alignment | not modelled | 11.2 | 39 | **Fold:**Domain of alpha and beta subunits of F1 ATP synthase-like<br>**Superfamily:**Aminopeptidase/glucanase lid domain<br>**Family:**Aminopeptidase/glucanase lid domain |
| 96 | d1pama3 | Alignment | not modelled | 11.2 | 18 | **Fold:**Glycosyl hydrolase domain<br>**Superfamily:**Glycosyl hydrolase domain<br>**Family:**alpha-Amylases, C-terminal beta-sheet domain |
| 97 | c6a4zA_ | Alignment | not modelled | 11.0 | 7 | **PDB header:**biosynthetic protein<br>**Chain:** A: **PDB Molecule:**chap protein;<br>**PDBTitle:** oxidase chap |
| 98 | c4qb5A_ | Alignment | not modelled | 11.0 | 14 | **PDB header:**oxidoreductase<br>**Chain:** A: **PDB Molecule:**glyoxalase/bleomycin resistance protein/dioxygenase;<br>**PDBTitle:** crystal structure of a glyoxalase/bleomycin resistance protein from2 albidiferax ferrireducens t118 |
| 99 | c4n04B_ | Alignment | not modelled | 11.0 | 17 | **PDB header:**oxidoreductase<br>**Chain:** B: **PDB Molecule:**glyoxalase/bleomycin resistance protein/dioxygenase;<br>**PDBTitle:** the crystal structure of glyoxalase / bleomycin resistance protein2 from catenulispora acidiphila dsm 44928 |
