## Supplementary material for "Nanopore long reads enable the first complete genome assembly of a Malaysian *Vibrio parahaemolyticus* isolate bearing the pVa plasmid associated with acute hepatopancreatic necrosis disease": Supplmental File 2 (from another version of this preprint: version 1, 2019-12-02)

### Phyre2

| | |
|---|---|
| Email | |
| Description | Undefined |
| Date | Mon Oct 21 04:21:53 BST 2019 |
| Unique Job ID | 8d346f76ac781310 |

Detailed template information

| # | Template | Alignment Coverage | 3D Model | Confidence | % i.d. | Template Information |
|---|---|---|---|---|---|---|
| 1 | c1vw2H_ | Alignment | 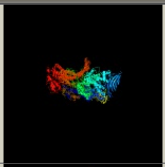 | 100.0 | 36 | **PDB header:**toxin<br>**Chain:** H: **PDB Molecule:**tcda1;<br>**PDBTitle:** crystal structure of tcda1 |
| 2 | c6ogdB_ | Alignment | 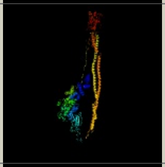 | 100.0 | 35 | **PDB header:**toxin<br>**Chain:** B: **PDB Molecule:**toxin subunit yena2;<br>**PDBTitle:** cryo-em structure of yentca in its prepore state |
| 3 | c6ogdD_ | Alignment | 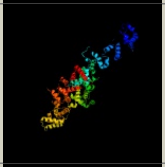 | 100.0 | 27 | **PDB header:**toxin<br>**Chain:** D: **PDB Molecule:**toxin subunit yena1;<br>**PDBTitle:** cryo-em structure of yentca in its prepore state |
| 4 | c2pohA_ | Alignment | 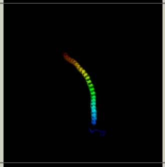 | 94.1 | 13 | **PDB header:**viral protein<br>**Chain:** A: **PDB Molecule:**head completion protein;<br>**PDBTitle:** structure of phage p22 tail needle gp26 |
| 5 | c3uuxD_ | Alignment | 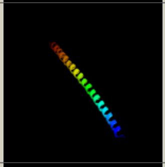 | 88.5 | 14 | **PDB header:**apoptosis<br>**Chain:** D: **PDB Molecule:**mitochondrial division protein 1;<br>**PDBTitle:** crystal structure of yeast fis1 in complex with mdv1 fragment2 containing n-terminal extension and coiled coil domains |
| 6 | c4rh7A_ | Alignment | 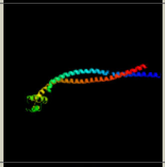 | 82.0 | 13 | **PDB header:**motor protein<br>**Chain:** A: **PDB Molecule:**green fluorescent protein/cytoplasmic dynein 2 heavy chain<br>**PDBTitle:** crystal structure of human cytoplasmic dynein 2 motor domain in2 complex with adp.vi |
| 7 | d1szpa1 | Alignment | 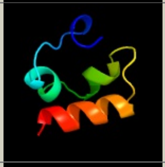 | 80.6 | 20 | **Fold:**SAM domain-like<br>**Superfamily:**Rad51 N-terminal domain-like<br>**Family:**DNA repair protein Rad51, N-terminal domain |
| 8 | c6e2jB_ | Alignment | 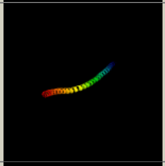 | 78.3 | 16 | **PDB header:**protein fibril<br>**Chain:** B: **PDB Molecule:**keratin, type i cytoskeletal 10;<br>**PDBTitle:** crystal structure of the heterocomplex between human keratin 1 coil 1b2 containing s233l mutation and wild-type human keratin 10 coil 1b |
| 9 | c5ew5C_ | Alignment | 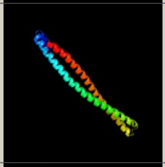 | 75.6 | 14 | **PDB header:**hydrolase<br>**Chain:** C: **PDB Molecule:**colicin-e9;<br>**PDBTitle:** crystal structure of colicin e9 in complex with its immunity protein2 im9 |
| 10 | c3vkgA_ | Alignment | 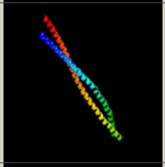 | 71.3 | 12 | **PDB header:**motor protein<br>**Chain:** A: **PDB Molecule:**dynein heavy chain, cytoplasmic;<br>**PDBTitle:** x-ray structure of an mtbd truncation mutant of dynein motor domain |
| 11 | c2xu6B_ | Alignment | 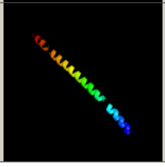 | 71.2 | 15 | **PDB header:**protein binding<br>**Chain:** B: **PDB Molecule:**mdv1 coiled coil;<br>**PDBTitle:** mdv1 coiled coil domain |

| | | | | | | |
|---|---|---|---|---|---|---|
| 12 | d1zeta2 | Alignment | 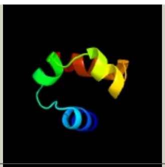 | 71.2 | 18 | **Fold:**DNA/RNA polymerases<br>**Superfamily:**DNA/RNA polymerases<br>**Family:**Lesion bypass DNA polymerase (Y-family), catalytic domain |
| 13 | c6a9pD_ | Alignment | 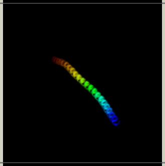 | 70.8 | 9 | **PDB header:**structural protein<br>**Chain:** D: **PDB Molecule:**glial fibrillary acidic protein;<br>**PDBTitle:** crystal structure of the human glial fibrillary acidic protein 1b2 domain |
| 14 | c1wx4B_ | Alignment | 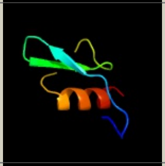 | 68.6 | 20 | **PDB header:**oxidoreductase/metal transport<br>**Chain:** B: **PDB Molecule:**melc;<br>**PDBTitle:** crystal structure of the oxy-form of the copper-bound streptomyces2 castaneoglobisporus tyrosinase complexed with a caddie protein3 prepared by the addition of dithiothreitol |
| 15 | d1b22a_ | Alignment | 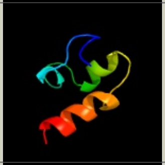 | 68.3 | 22 | **Fold:**SAM domain-like<br>**Superfamily:**Rad51 N-terminal domain-like<br>**Family:**DNA repair protein Rad51, N-terminal domain |
| 16 | c1b22A_ | Alignment | 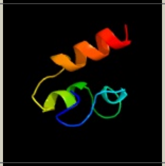 | 68.3 | 22 | **PDB header:**dna binding protein<br>**Chain:** A: **PDB Molecule:**dna repair protein rad51;<br>**PDBTitle:** rad51 (n-terminal domain) |
| 17 | c5j65A_ | Alignment | 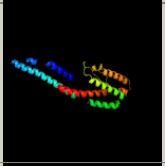 | 67.3 | 11 | **PDB header:**toxin<br>**Chain:** A: **PDB Molecule:**pesticidal crystal protein cry6aa;<br>**PDBTitle:** crystal structure of trypsin activated cry6aa |
| 18 | c3n4xB_ | Alignment | 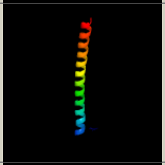 | 67.1 | 19 | **PDB header:**replication<br>**Chain:** B: **PDB Molecule:**monopolin complex subunit csm1;<br>**PDBTitle:** structure of csm1 full-length |
| 19 | c5mqrA_ | Alignment | 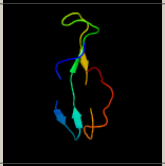 | 67.0 | 16 | **PDB header:**hydrolase<br>**Chain:** A: **PDB Molecule:**beta-l-arabinobiosidase;<br>**PDBTitle:** sialidase bt_1020 |
| 20 | c2hpcF_ | Alignment | 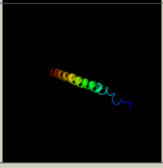 | 66.9 | 11 | **PDB header:**blood clotting<br>**Chain:** F: **PDB Molecule:**fibrinogen, gamma polypeptide;<br>**PDBTitle:** crystal structure of fragment d from human fibrinogen complexed with2 gly-pro-arg-pro-amide. |
| 21 | d2i1qa1 | Alignment | not modelled | 66.6 | 21 | **Fold:**SAM domain-like<br>**Superfamily:**Rad51 N-terminal domain-like<br>**Family:**DNA repair protein Rad51, N-terminal domain |
| 22 | d1pzna1 | Alignment | not modelled | 65.8 | 27 | **Fold:**SAM domain-like<br>**Superfamily:**Rad51 N-terminal domain-like<br>**Family:**DNA repair protein Rad51, N-terminal domain |
| 23 | c3lsoA_ | Alignment | not modelled | 65.7 | 31 | **PDB header:**membrane protein<br>**Chain:** A: **PDB Molecule:**putative membrane anchored protein;<br>**PDBTitle:** crystal structure of putative membrane anchored protein from2 corynebacterium diphtheriae |
| 24 | c5d3aA_ | Alignment | not modelled | 62.0 | 14 | **PDB header:**motor protein<br>**Chain:** A: **PDB Molecule:**kinesin-like protein kif21a;<br>**PDBTitle:** kif21a regulatory coiled coil |
| 25 | c3vkhD_ | Alignment | not modelled | 61.6 | 9 | **PDB header:**motor protein<br>**Chain:** D: **PDB Molecule:**<br>**PDBTitle:** x-ray structure of a functional full-length dynein motor domain |
| 26 | c3swkB_ | Alignment | not modelled | 60.9 | 12 | **PDB header:**structural protein<br>**Chain:** B: **PDB Molecule:**vimentin;<br>**PDBTitle:** crystal structure of vimentin coil1b fragment |
| 27 | c3m9bK_ | Alignment | not modelled | 55.6 | 23 | **PDB header:**chaperone<br>**Chain:** K: **PDB Molecule:**proteasome-associated atpase;<br>**PDBTitle:** crystal structure of the amino terminal coiled coil domain and the2 inter domain of the mycobacterium tuberculosis proteasomal atpase mpa |
| 28 | d1szpb1 | Alignment | not modelled | 54.9 | 23 | **Fold:**SAM domain-like<br>**Superfamily:**Rad51 N-terminal domain-like<br>**Family:**DNA repair protein Rad51, N-terminal domain |
| | | | | | | **Fold:**STAT-like |

| | | | | | | |
|---|---|---|---|---|---|---|
| 29 | d1tkna_ | Alignment | not modelled | 54.3 | 10 | **Superfamily:**CAPPD, an extracellular domain of amyloid beta A4 protein<br>**Family:**CAPPD, an extracellular domain of amyloid beta A4 protein |
| 30 | d1nera_ | Alignment | not modelled | 54.3 | 13 | **Fold:**lambda repressor-like DNA-binding domains<br>**Superfamily:**lambda repressor-like DNA-binding domains<br>**Family:**Phage repressors |
| 31 | c6ec0A_ | Alignment | not modelled | 52.5 | 12 | **PDB header:**protein fibril<br>**Chain:** A: **PDB Molecule:**keratin 1;<br>**PDBTitle:** crystal structure of the wild-type heterocomplex between coil 1b2 domains of human intermediate filament proteins keratin 1 (krt1) and3 keratin 10 (krt10) |
| 32 | c4dezA_ | Alignment | not modelled | 51.0 | 18 | **PDB header:**transferase<br>**Chain:** A: **PDB Molecule:**dna polymerase iv 1;<br>**PDBTitle:** structure of msdpo4 |
| 33 | c3pmrB_ | Alignment | not modelled | 50.7 | 14 | **PDB header:**cell adhesion<br>**Chain:** B: **PDB Molecule:**amyloid-like protein 1;<br>**PDBTitle:** crystal structure of e2 domain of human amyloid precursor-like protein2 1 |
| 34 | c6iu3A_ | Alignment | not modelled | 50.2 | 10 | **PDB header:**metal transport<br>**Chain:** A: **PDB Molecule:**vit1;<br>**PDBTitle:** crystal structure of iron transporter vit1 with zinc ions |
| 35 | c3mr2A_ | Alignment | not modelled | 49.0 | 13 | **PDB header:**transferase/dna<br>**Chain:** A: **PDB Molecule:**dna polymerase eta;<br>**PDBTitle:** human dna polymerase eta in complex with normal dna and incoming2 nucleotide (nrm) |
| 36 | c2fllA_ | Alignment | not modelled | 47.9 | 18 | **PDB header:**replication/dna<br>**Chain:** A: **PDB Molecule:**dna polymerase iota;<br>**PDBTitle:** ternary complex of human dna polymerase iota with dna and dttp |
| 37 | c6basA_ | Alignment | not modelled | 47.9 | 22 | **PDB header:**transferase<br>**Chain:** A: **PDB Molecule:**peptidoglycan glycosyltransferase roda;<br>**PDBTitle:** crystal structure of thermus thermophilus rod shape determining2 protein roda d255a mutant (q5six3_thet8) |
| 38 | d1jx4a2 | Alignment | not modelled | 47.2 | 22 | **Fold:**DNA/RNA polymerases<br>**Superfamily:**DNA/RNA polymerases<br>**Family:**Lesion bypass DNA polymerase (Y-family), catalytic domain |
| 39 | c1deqO_ | Alignment | not modelled | 46.2 | 10 | **PDB header:**blood clotting<br>**Chain:** O: **PDB Molecule:**fibrinogen (beta chain);<br>**PDBTitle:** the crystal structure of modified bovine fibrinogen (at ~42 angstrom resolution) |
| 40 | c4r0rA_ | Alignment | not modelled | 45.6 | 16 | **PDB header:**biosynthetic protein<br>**Chain:** A: **PDB Molecule:**eboizn21;<br>**PDBTitle:** ebolavirus gp prehairpin intermediate mimic |
| 41 | c1t3nB_ | Alignment | not modelled | 43.7 | 20 | **PDB header:**replication/dna<br>**Chain:** B: **PDB Molecule:**polymerase (dna directed) iota;<br>**PDBTitle:** structure of the catalytic core of dna polymerase iota in complex with2 dna and dttp |
| 42 | c5ldwV_ | Alignment | not modelled | 43.6 | 12 | **PDB header:**oxidoreductase<br>**Chain:** V: **PDB Molecule:**nadh dehydrogenase [ubiquinone] 1 alpha subcomplex subunit<br>**PDBTitle:** structure of mammalian respiratory complex i, class1 |
| 43 | c4l9uB_ | Alignment | not modelled | 43.6 | 21 | **PDB header:**signaling protein<br>**Chain:** B: **PDB Molecule:**ras guanyl-releasing protein 1;<br>**PDBTitle:** structure of c-terminal coiled coil of rasgrp1 |
| 44 | c5mg8B_ | Alignment | not modelled | 40.1 | 18 | **PDB header:**recombination<br>**Chain:** B: **PDB Molecule:**structural maintenance of chromosomes protein 6;<br>**PDBTitle:** crystal structure of the s.pombe smc5/6 hinge domain |
| 45 | d1rw6a_ | Alignment | not modelled | 40.1 | 10 | **Fold:**STAT-like<br>**Superfamily:**CAPPD, an extracellular domain of amyloid beta A4 protein<br>**Family:**CAPPD, an extracellular domain of amyloid beta A4 protein |
| 46 | c5gnaB_ | Alignment | not modelled | 36.4 | 16 | **PDB header:**gene regulation<br>**Chain:** B: **PDB Molecule:**flagellar hook-associated protein 2;<br>**PDBTitle:** crystal structure of flagellin assembly related protein |
| 47 | d1k1sa2 | Alignment | not modelled | 36.1 | 16 | **Fold:**DNA/RNA polymerases<br>**Superfamily:**DNA/RNA polymerases<br>**Family:**Lesion bypass DNA polymerase (Y-family), catalytic domain |
| 48 | c5tptA_ | Alignment | not modelled | 35.7 | 12 | **PDB header:**protein fibril<br>**Chain:** A: **PDB Molecule:**amyloid-like protein 2;<br>**PDBTitle:** the crystal structure of amyloid precursor-like protein 2 (aplp2) e22 domain |
| 49 | c3gqcB_ | Alignment | not modelled | 35.3 | 16 | **PDB header:**transferase/dna<br>**Chain:** B: **PDB Molecule:**dna repair protein rev1;<br>**PDBTitle:** structure of human rev1-dna-dntp ternary complex |
| 50 | c3vkgB_ | Alignment | not modelled | 34.7 | 11 | **PDB header:**motor protein<br>**Chain:** B: **PDB Molecule:**dynein heavy chain, cytoplasmic;<br>**PDBTitle:** x-ray structure of an mtbd truncation mutant of dynein motor domain |
| 51 | c1yodB_ | Alignment | not modelled | 34.3 | 100 | **PDB header:**de novo protein<br>**Chain:** B: **PDB Molecule:**water-solublized phospholamban;<br>**PDBTitle:** crystal structure of a water soluble analog of phospholamban |
| 52 | c5jvpC_ | Alignment | not modelled | 34.2 | 25 | **PDB header:**motor protein<br>**Chain:** C: **PDB Molecule:**chimera protein of centromere-associated protein e and<br>**PDBTitle:** the neck-linker and alpha 7 helix of homo sapiens cenp-e |
| 53 | c5hx0A_ | Alignment | not modelled | 33.6 | 18 | **PDB header:**structural genomics, unknown function<br>**Chain:** A: **PDB Molecule:**uncharacterized protein dfer_1899;<br>**PDBTitle:** crystal structure of unknown function protein dfer_18992 fromdyadobacter fermentans dsm 18053 |
| 54 | c2kr0A_ | Alignment | not modelled | 33.4 | 14 | **PDB header:**protein binding<br>**Chain:** A: **PDB Molecule:**proteasomal ubiquitin receptor adrm1;<br>**PDBTitle:** solution structure of the proteasome ubiquitin receptor rpn13 |
| [illegible] | [illegible] | [illegible] | [illegible] | [illegible] | [illegible] | **PDB header:**de novo protein<br>[illegible] |

| | | | | | | |
|---|---|---|---|---|---|---|
| 55 | c1yodA_ | Alignment | not modelled | 33.3 | 100 | **Chain:** A: **PDB Molecule:**water-solubilized phospholamban; **PDBTitle:** crystal structure of a water soluble analog of phospholamban |
| 56 | c5m9eA_ | Alignment | not modelled | 33.0 | 17 | **PDB header:**cell cycle **Chain:** A: **PDB Molecule:**microtubule integrity protein mal3; **PDBTitle:** interactions between the mal3 eb1-like domain and dis1 |
| 57 | c6ppbW_ | Alignment | not modelled | 33.0 | 14 | **PDB header:**viral protein **Chain:** W: **PDB Molecule:**major capsid protein; **PDBTitle:** kaposi's sarcoma-associated herpesvirus (kshv), c5 portal vertex2 structure |
| 58 | c4ir1A_ | Alignment | not modelled | 31.7 | 13 | **PDB header:**transferase/dna **Chain:** A: **PDB Molecule:**dna polymerase iv; **PDBTitle:** polymerase-dna complex |
| 59 | c5oqmh_ | Alignment | not modelled | 31.4 | 18 | **PDB header:**transcription **Chain:** H: **PDB Molecule:**dna-directed rna polymerases i, ii, and iii subunit rpabc3; **PDBTitle:** structure of yeast transcription pre-initiation complex with tfiih and2 core mediator |
| 60 | c1k1qA_ | Alignment | not modelled | 31.3 | 16 | **PDB header:**transcription **Chain:** A: **PDB Molecule:**dbh protein; **PDBTitle:** crystal structure of a dinb family error prone dna2 polymerase from sulfolobus solfataricus |
| 61 | c4v19J_ | Alignment | not modelled | 30.6 | 24 | **PDB header:**ribosome **Chain:** J: **PDB Molecule:**mitoribosomal protein ul10m, mrpl10; **PDBTitle:** structure of the large subunit of the mammalian mitoribosome, part 12 of 2 |
| 62 | c4p3gC_ | Alignment | not modelled | 30.6 | 19 | **PDB header:**rna binding protein **Chain:** C: **PDB Molecule:**signal recognition particle subunit srp68; **PDBTitle:** structure of the srp68-rbd from chaetomium thermophilum |
| 63 | c4l9uA_ | Alignment | not modelled | 30.1 | 23 | **PDB header:**signaling protein **Chain:** A: **PDB Molecule:**ras guanyl-releasing protein 1; **PDBTitle:** structure of c-terminal coiled coil of rasgrp1 |
| 64 | c5x90B_ | Alignment | not modelled | 30.0 | 47 | **PDB header:**protein transport **Chain:** B: **PDB Molecule:**icmw; **PDBTitle:** structure of dotl(656-783)-icms-icmw-lvga derived from legionella2 pneumophila |
| 65 | c5cqvB_ | Alignment | not modelled | 30.0 | 17 | **PDB header:**unknown function **Chain:** B: **PDB Molecule:**uncharacterized protein; **PDBTitle:** crystal structure of uncharacterized protein q8dwv2 from streptococcus2 agalactiae |
| 66 | d2p12a1 | Alignment | not modelled | 29.8 | 21 | **Fold:**FomD barrel-like **Superfamily:**FomD-like **Family:**FomD-like |
| 67 | c3a0oB_ | Alignment | not modelled | 29.8 | 12 | **PDB header:**lyase **Chain:** B: **PDB Molecule:**oligo alginate lyase; **PDBTitle:** crystal structure of alginate lyase from agrobacterium tumefaciens c58 |
| 68 | c4e61A_ | Alignment | not modelled | 28.8 | 19 | **PDB header:**cell cycle **Chain:** A: **PDB Molecule:**protein bim1; **PDBTitle:** crystal structure of the eb1-like motif of bim1p |
| 69 | d1dvoa_ | Alignment | not modelled | 28.2 | 14 | **Fold:**FinO-like **Superfamily:**FinO-like **Family:**FinO-like |
| 70 | c1s97D_ | Alignment | not modelled | 28.0 | 21 | **PDB header:**transferase/dna **Chain:** D: **PDB Molecule:**dna polymerase iv; **PDBTitle:** dpo4 with gt mismatch |
| 71 | c5wjbA_ | Alignment | not modelled | 27.3 | 12 | **PDB header:**actin/dna binding protein **Chain:** A: **PDB Molecule:**capsid assembly scaffolding protein,myosin-7; **PDBTitle:** crystal structure of amino acids 1733-1797 of human beta cardiac2 myosin fused to gp7 |
| 72 | c5cwsJ_ | Alignment | not modelled | 27.1 | 9 | **PDB header:**protein transport **Chain:** J: **PDB Molecule:**nucleoporin nup49; **PDBTitle:** crystal structure of the intact chaetomium thermophilum nsp1-nup49-2 nup57 channel nucleoporin heterotrimer bound to its nic96 nuclear3 pore complex attachment site |
| 73 | c1pl5A_ | Alignment | not modelled | 26.9 | 12 | **PDB header:**dna binding protein/transcription **Chain:** A: **PDB Molecule:**regulatory protein sir4; **PDBTitle:** crystal structure analysis of the sir4p c-terminal coiled2 coil |
| 74 | c4gvsA_ | Alignment | not modelled | 26.8 | 9 | **PDB header:**hydrolase **Chain:** A: **PDB Molecule:**methenyltetrahydromethanopterin cyclohydrolase; **PDBTitle:** x-ray structure of the archaeoglobus fulgidus methenyl-2 tetrahydromethanopterin cyclohydrolase in complex with n5-formyl-3 tetrahydromethanopterin |
| 75 | c1nyhA_ | Alignment | not modelled | 26.8 | 12 | **PDB header:**transcription repressor **Chain:** A: **PDB Molecule:**regulatory protein sir4; **PDBTitle:** crystal structure of the coiled-coil dimerization motif of sir4 |
| 76 | c2rf0D_ | Alignment | not modelled | 26.7 | 27 | **PDB header:**transferase **Chain:** D: **PDB Molecule:**mitogen-activated protein kinase kinase kinase 10; **PDBTitle:** crystal structure of human mixed lineage kinase map3k10 sh3 domain |
| 77 | c1ei3E_ | Alignment | not modelled | 26.7 | 6 | **PDB header:**blood clotting **Chain:** E: **PDB Molecule:**fibrinogen; **PDBTitle:** crystal structure of native chicken fibrinogen |
| 78 | c2kc1A_ | Alignment | not modelled | 26.6 | 21 | **PDB header:**structural protein **Chain:** A: **PDB Molecule:**mkiaa1027 protein; **PDBTitle:** nmr structure of the f0 domain (residues 0-85) of the talin2 ferm domain |
| 79 | c2cblA_ | Alignment | not modelled | 26.5 | 23 | **PDB header:**complex (proto-oncogene/peptide) **Chain:** A: **PDB Molecule:**proto-oncogene cbl; **PDBTitle:** n-terminal domain of cbl in complex with its binding site2 on zap-70 |
| | | | | | | **PDB header:**contractile protein |

| | | | | | | |
|---|---|---|---|---|---|---|
| 80 | c3u1aC_ | Alignment | not modelled | 26.4 | 12 | **Chain:** C: **PDB Molecule:**smooth muscle tropomyosin alpha; **PDBTitle:** n-terminal 81-aa fragment of smooth muscle tropomyosin alpha |
| 81 | c3c1dA_ | Alignment | not modelled | 26.1 | 19 | **PDB header:**recombination, dna binding protein **Chain:** A: **PDB Molecule:**regulatory protein recx; **PDBTitle:** x-ray crystal structure of recx |
| 82 | c6cl6E_ | Alignment | not modelled | 25.9 | 22 | **PDB header:**viral protein **Chain:** E: **PDB Molecule:**tail fiber protein; **PDBTitle:** structure of p. aeruginosa r2 pyocin fiber pa0620 comprising c-2 terminal residues 323-691 |
| 83 | c6girA_ | Alignment | not modelled | 25.4 | 9 | **PDB header:**cytosolic protein **Chain:** A: **PDB Molecule:**serine--trna ligase, cytoplasmic; **PDBTitle:** arabidopsis thaliana cytosolic seryl-trna synthetase |
| 84 | c6gy6Q_ | Alignment | not modelled | 25.2 | 14 | **PDB header:**toxin **Chain:** Q: **PDB Molecule:**xaxa; **PDBTitle:** xaxab pore complex from xenorhabdus nematophila |
| 85 | d3buxb2 | Alignment | not modelled | 25.1 | 23 | **Fold:**N-cbl like **Superfamily:**N-terminal domain of cbl (N-cbl) **Family:**N-terminal domain of cbl (N-cbl) |
| 86 | d1t94a2 | Alignment | not modelled | 24.5 | 8 | **Fold:**DNA/RNA polymerases **Superfamily:**DNA/RNA polymerases **Family:**Lesion bypass DNA polymerase (Y-family), catalytic domain |
| 87 | c6g3eA_ | Alignment | not modelled | 24.1 | 10 | **PDB header:**lyase **Chain:** A: **PDB Molecule:**argininosuccinate lyase; **PDBTitle:** crystal structure of edds lyase in complex with formate |
| 88 | d1lb2b_ | Alignment | not modelled | 24.1 | 8 | **Fold:**SAM domain-like **Superfamily:**C-terminal domain of RNA polymerase alpha subunit **Family:**C-terminal domain of RNA polymerase alpha subunit |
| 89 | c5gasN_ | Alignment | not modelled | 23.8 | 9 | **PDB header:**hydrolase **Chain:** N: **PDB Molecule:**archaeal/vacuolar-type h+-atpase subunit i; **PDBTitle:** thermus thermophilus v/a-atpase, conformation 2 |
| 90 | c6o7xa_ | Alignment | not modelled | 23.6 | 4 | **PDB header:**membrane protein **Chain:** A: **PDB Molecule:**vacuolar atp synthase catalytic subunit a; **PDBTitle:** saccharomyces cerevisiae v-atpase stv1-v1vo state 3 |
| 91 | c6ppdA_ | Alignment | not modelled | 23.4 | 24 | **PDB header:**viral protein **Chain:** A: **PDB Molecule:**small capsomere-interacting protein; **PDBTitle:** kaposi's sarcoma-associated herpesvirus (kshv), c1 penton vertex2 register, catc-absent structure |
| 92 | c4xwhA_ | Alignment | not modelled | 23.2 | 14 | **PDB header:**hydrolase **Chain:** A: **PDB Molecule:**alpha-n-acetylglucosaminidase; **PDBTitle:** crystal structure of the human n-acetyl-alpha-glucosaminidase |
| 93 | d2j07a2 | Alignment | not modelled | 22.3 | 29 | **Fold:**Cryptochrome/photolyase, N-terminal domain **Superfamily:**Cryptochrome/photolyase, N-terminal domain **Family:**Cryptochrome/photolyase, N-terminal domain |
| 94 | c3o0lB_ | Alignment | not modelled | 22.2 | 23 | **PDB header:**structural genomics, unknown function **Chain:** B: **PDB Molecule:**uncharacterized protein; **PDBTitle:** crystal structure of a pfam duf1425 family member (shew_1734) from2 shewanella sp. pv-4 at 1.81 a resolution |
| 95 | d1y57a1 | Alignment | not modelled | 22.0 | 33 | **Fold:**SH3-like barrel **Superfamily:**SH3-domain **Family:**SH3-domain |
| 96 | c2lq4p_ | Alignment | not modelled | 22.0 | 40 | **PDB header:**de novo protein **Chain:** P: **PDB Molecule:** **PDBTitle:** structural characterization of an lpa1 second extracellular loop2 mimetic with a self-assembling coiled-coil folding constraint |
| 97 | c6u94A_ | Alignment | not modelled | 21.9 | 11 | **PDB header:**lipid transport **Chain:** A: **PDB Molecule:**rnd efflux system, outer membrane lipoprotein, nodt family; **PDBTitle:** structure of rnd efflux system, outer membrane lipoprotein, nodt2 family from burkholderia mallei atcc 23344 |
| 98 | c2qv6D_ | Alignment | not modelled | 21.8 | 17 | **PDB header:**hydrolase **Chain:** D: **PDB Molecule:**gtp cyclohydrolase iii; **PDBTitle:** gtp cyclohydrolase iii from m. jannaschii (mj0145)2 complexed with gtp and metal ions |
| 99 | c5lskA_ | Alignment | not modelled | 21.6 | 19 | **PDB header:**cell cycle **Chain:** A: **PDB Molecule:**protein mis12 homolog; **PDBTitle:** crystal structure of the human kinetochore mis12-cenp-c complex |
| 100 | c6orjA_ | Alignment | not modelled | 21.4 | 27 | **PDB header:**viral protein **Chain:** A: **PDB Molecule:**phikz164; **PDBTitle:** central spike of phikz phage tail |
| 101 | c3bunB_ | Alignment | not modelled | 21.3 | 23 | **PDB header:**ligase/signaling protein **Chain:** B: **PDB Molecule:**e3 ubiquitin-protein ligase cbl; **PDBTitle:** crystal structure of c-cbl-tkb domain complexed with its binding motif2 in sprouty4 |
| 102 | d2h1ta1 | Alignment | not modelled | 21.2 | 26 | **Fold:**Spiral beta-roll **Superfamily:**PA1994-like **Family:**PA1994-like |
| 103 | c4uirB_ | Alignment | not modelled | 21.1 | 17 | **PDB header:**lyase **Chain:** B: **PDB Molecule:**oleate hydratase; **PDBTitle:** structure of oleate hydratase from elizabethkingia2 meningoseptica |
| 104 | d1z3eb1 | Alignment | not modelled | 21.1 | 14 | **Fold:**SAM domain-like **Superfamily:**C-terminal domain of RNA polymerase alpha subunit **Family:**C-terminal domain of RNA polymerase alpha subunit |
| 105 | c3ol1A_ | Alignment | not modelled | 21.0 | 12 | **PDB header:**structural protein **Chain:** A: **PDB Molecule:**vimentin; **PDBTitle:** crystal structure of vimentin (fragment 144-251) from homo sapiens,2 northeast structural genomics consortium target hr4796b |
| | | | | | | **PDB header:**contractile protein |

| | | | | | | |
|---|---|---|---|---|---|---|
| 106 | c1zuyB_ | Alignment | not modelled | 20.9 | 36 | **Chain:** B: **PDB Molecule:**myosin-5 isoform;<br>**PDBTitle:** high-resolution structure of yeast myo5 sh3 domain |
| 107 | c4f4hA_ | Alignment | not modelled | 20.9 | 15 | **PDB header:**ligase<br>**Chain:** A: **PDB Molecule:**glutamine dependent nad+ synthetase;<br>**PDBTitle:** crystal structure of a glutamine dependent nad+ synthetase from2 burkholderia thailandensis |
| 108 | c1o7dC_ | Alignment | not modelled | 20.5 | 13 | **PDB header:**hydrolase<br>**Chain:** C: **PDB Molecule:**lysosomal alpha-mannosidase;<br>**PDBTitle:** the structure of the bovine lysosomal a-mannosidase suggests a novel2 mechanism for low ph activation |
| 109 | c4xghA_ | Alignment | not modelled | 20.1 | 27 | **PDB header:**transferase<br>**Chain:** A: **PDB Molecule:**citrate synthase;<br>**PDBTitle:** x-ray crystal structure of citrate synthase from burkholderia2 thailandensis |
