## Supplementary material for "Nanopore long reads enable the first complete genome assembly of a Malaysian *Vibrio parahaemolyticus* isolate bearing the pVa plasmid associated with acute hepatopancreatic necrosis disease": Supplmental File 3 (from another version of this preprint: version 1, 2019-12-02)

### Phyre2


| | |
|---|---|
| Email | |
| Description | tccc____ |
| Date | Fri Nov 8 00:06:32 GMT 2019 |
| Unique Job ID | b0d47cdf2c0b1591 |


Detailed template information

| # | Template | Alignment Coverage | 3D Model | Confidence | % i.d. | Template Information |
|---|---|---|---|---|---|---|
| 1 | c5kisB_ | Alignment | 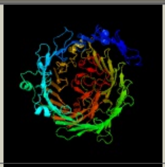 | 100.0 | 72 | **PDB header:**toxin **Chain:** B: **PDB Molecule:**rhs2; **PDBTitle:** yenb/rhs2 complex |
| 2 | c4o9xA_ | Alignment | 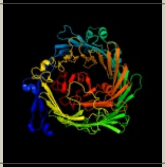 | 100.0 | 59 | **PDB header:**toxin **Chain:** A: **PDB Molecule:**tcdb2, tccc3; **PDBTitle:** crystal structure of tcdb2-tccc3 |
| 3 | c4iglB_ | Alignment | 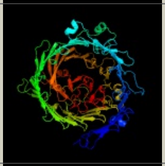 | 100.0 | 65 | **PDB header:**toxin **Chain:** B: **PDB Molecule:**yenc2; **PDBTitle:** structure of the rhs-repeat containing bc component of the secreted2 abc toxin complex from yersinia entomophaga |
| 4 | c6fayA_ | Alignment | 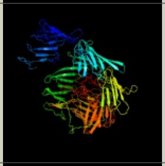 | 100.0 | 16 | **PDB header:**cell adhesion **Chain:** A: **PDB Molecule:**odz3 protein; **PDBTitle:** teneurin3 monomer |
| 5 | c6fb3A_ | Alignment | 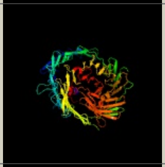 | 100.0 | 15 | **PDB header:**cell adhesion **Chain:** A: **PDB Molecule:**teneurin-2; **PDBTitle:** teneurin 2 partial extracellular domain |
| 6 | c6cmxA_ | Alignment | 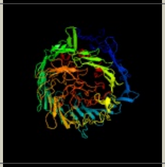 | 100.0 | 15 | **PDB header:**membrane protein **Chain:** A: **PDB Molecule:**teneurin-2; **PDBTitle:** human teneurin 2 extra-cellular region |
| 7 | c4iglA_ | Alignment | 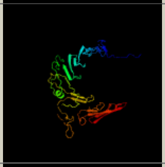 | 99.9 | 13 | **PDB header:**toxin **Chain:** A: **PDB Molecule:**yenb; **PDBTitle:** structure of the rhs-repeat containing bc component of the secreted2 abc toxin complex from yersinia entomophaga |
| 8 | c4ghbA_ | Alignment | 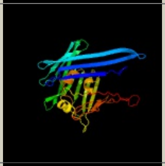 | 99.8 | 11 | **PDB header:**structural genomics, unknown function **Chain:** A: **PDB Molecule:**hypothetical protein; **PDBTitle:** crystal structure of a porin-like protein (bacuni_01323) from2 bacteroides uniformis atcc 8492 at 2.32 a resolution |
| 9 | c2oy7A_ | Alignment | 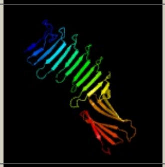 | 80.1 | 11 | **PDB header:**membrane protein **Chain:** A: **PDB Molecule:**outer surface protein a; **PDBTitle:** the crystal structure of ospa mutant |
| 10 | d1x3za1 | Alignment | 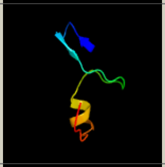 | 57.0 | 20 | **Fold:**Cysteine proteinases **Superfamily:**Cysteine proteinases **Family:**Transglutaminase core |
| 11 | d2f4ma1 | Alignment | 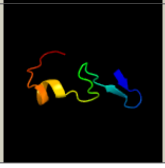 | 45.9 | 26 | **Fold:**Cysteine proteinases **Superfamily:**Cysteine proteinases **Family:**Transglutaminase core |

| | | | | | | |
|---|---|---|---|---|---|---|
| 12 | c5lo2A_ | Alignment | 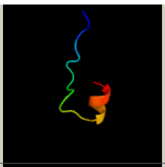 | 44.9 | 40 | **PDB header:**structural protein<br>**Chain:** A: **PDB Molecule:**ppatyr;<br>**PDBTitle:** engineering protein stability with atomic precision in a monomeric2 miniprotein |
| 13 | c2mxwA_ | Alignment | 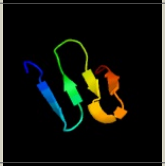 | 41.9 | 41 | **PDB header:**unknown function<br>**Chain:** A: **PDB Molecule:**rna-binding protein 10;<br>**PDBTitle:** solution nmr structure of the ocre domain of rbm10 |
| 14 | c3a0mF_ | Alignment | 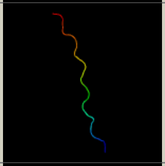 | 31.9 | 57 | **PDB header:**structural protein<br>**Chain:** F: **PDB Molecule:**collagen-like peptide;<br>**PDBTitle:** structure of (ppg)4-ovg-(ppg)4, monoclinic, twinned crystal |
| 15 | c2nqoB_ | Alignment | 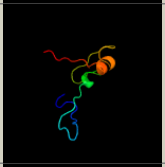 | 30.6 | 21 | **PDB header:**transferase<br>**Chain:** B: **PDB Molecule:**gamma-glutamyltranspeptidase;<br>**PDBTitle:** crystal structure of helicobacter pylori gamma-glutamyltranspeptidase |
| 16 | c3eswA_ | Alignment | 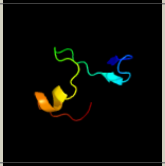 | 28.6 | 20 | **PDB header:**hydrolase<br>**Chain:** A: **PDB Molecule:**peptide-n(4)-(n-acetyl-beta-glucosaminyl)asparagine<br>**PDBTitle:** complex of yeast pngase with glcnac2-iac. |
| 17 | c3a19B_ | Alignment | 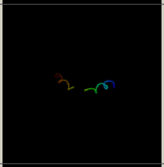 | 26.3 | 57 | **PDB header:**structural protein<br>**Chain:** B: **PDB Molecule:**collagen-like peptide;<br>**PDBTitle:** structure of (ppg)4-oog-(ppg)4_h monoclinic, twinned crystal |
| 18 | c2d3hA_ | Alignment | 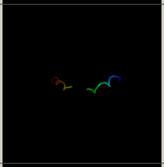 | 26.3 | 57 | **PDB header:**structural protein<br>**Chain:** A: **PDB Molecule:**collagen model peptides (pro-pro-gly)4-hyp-hyp-<br>**PDBTitle:** crystal structures of collagen model peptides (pro-pro-gly)2 4-hyp-hyp-gly-(pro-pro-gly)4 |
| 19 | c2d3hB_ | Alignment | 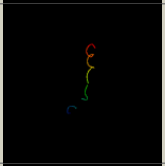 | 26.3 | 57 | **PDB header:**structural protein<br>**Chain:** B: **PDB Molecule:**collagen model peptides (pro-pro-gly)4-hyp-hyp-<br>**PDBTitle:** crystal structures of collagen model peptides (pro-pro-gly)2 4-hyp-hyp-gly-(pro-pro-gly)4 |
| 20 | c3wknG_ | Alignment | 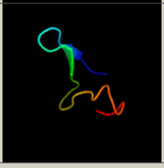 | 25.9 | 36 | **PDB header:**immune system<br>**Chain:** G: **PDB Molecule:**affinger p17;<br>**PDBTitle:** crystal structure of the artificial protein affinger p17 (af.p17)2 complexed with fc fragment of human igg |
| 21 | c5lo3A_ | Alignment | not modelled | 22.0 | 37 | **PDB header:**structural protein<br>**Chain:** A: **PDB Molecule:**ppaome;<br>**PDBTitle:** engineering protein stability with atomic precision in a monomeric2 miniprotein |
| 22 | c5lo4A_ | Alignment | not modelled | 22.0 | 37 | **PDB header:**structural protein<br>**Chain:** A: **PDB Molecule:**ppa-ch3;<br>**PDBTitle:** engineering protein stability with atomic precision in a monomeric2 miniprotein |
| 23 | c3a08C_ | Alignment | not modelled | 22.0 | 58 | **PDB header:**structural protein<br>**Chain:** C: **PDB Molecule:**collagen-like peptide;<br>**PDBTitle:** structure of (ppg)4-oog-(ppg)4, monoclinic, twinned crystal |
| 24 | c2d3hC_ | Alignment | not modelled | 21.6 | 57 | **PDB header:**structural protein<br>**Chain:** C: **PDB Molecule:**collagen model peptides (pro-pro-gly)4-hyp-hyp-<br>**PDBTitle:** crystal structures of collagen model peptides (pro-pro-gly)2 4-hyp-hyp-gly-(pro-pro-gly)4 |
| 25 | c3a19A_ | Alignment | not modelled | 21.6 | 57 | **PDB header:**structural protein<br>**Chain:** A: **PDB Molecule:**collagen-like peptide;<br>**PDBTitle:** structure of (ppg)4-oog-(ppg)4_h monoclinic, twinned crystal |
| 26 | c2d3hE_ | Alignment | not modelled | 21.6 | 57 | **PDB header:**structural protein<br>**Chain:** E: **PDB Molecule:**collagen model peptides (pro-pro-gly)4-hyp-hyp-<br>**PDBTitle:** crystal structures of collagen model peptides (pro-pro-gly)2 4-hyp-hyp-gly-(pro-pro-gly)4 |
| 27 | c3ah9D_ | Alignment | not modelled | 20.5 | 57 | **PDB header:**structural protein<br>**Chain:** D: **PDB Molecule:**collagen-like peptide;<br>**PDBTitle:** crystal structure of (pro-pro-gly)9 at 1.1 a resolution |
| 28 | c3a19D_ | Alignment | not modelled | 19.8 | 57 | **PDB header:**structural protein<br>**Chain:** D: **PDB Molecule:**collagen-like peptide;<br>**PDBTitle:** structure of (ppg)4-oog-(ppg)4_h monoclinic, twinned |

| | | | | | | |
|---|---|---|---|---|---|---|
| | | | | | | crystal |
| 29 | c3a08F_ | Alignment | not modelled | 19.6 | 56 | **PDB header:**structural protein<br>**Chain:** F: **PDB Molecule:**collagen-like peptide;<br>**PDBTitle:** structure of (ppg)4-oog-(ppg)4, monoclinic, twinned crystal |
| 30 | c1tz5A_ | Alignment | not modelled | 19.0 | 32 | **PDB header:**hormone/growth factor<br>**Chain:** A: **PDB Molecule:**chimera of pancreatic hormone and neuropeptide y;<br>**PDBTitle:** [pnpy19-23]-hpp bound to dpc micelles |
| 31 | c2vsaA_ | Alignment | not modelled | 18.9 | 9 | **PDB header:**toxin<br>**Chain:** A: **PDB Molecule:**mosquitocidal toxin;<br>**PDBTitle:** structure and mode of action of a mosquitocidal holotoxin |
| 32 | c5akoD_ | Alignment | not modelled | 18.5 | 40 | **PDB header:**unknown function<br>**Chain:** D: **PDB Molecule:**tse2;<br>**PDBTitle:** the complex of tse2 and tsi2 from pseudomonas aeruginosa |
| 33 | d1bcoa1 | Alignment | not modelled | 18.1 | 17 | **Fold:**mu transposase, C-terminal domain<br>**Superfamily:**mu transposase, C-terminal domain<br>**Family:**mu transposase, C-terminal domain |
| 34 | c2d3hF_ | Alignment | not modelled | 18.0 | 58 | **PDB header:**structural protein<br>**Chain:** F: **PDB Molecule:**collagen model peptides (pro-pro-gly)4-hyp-hyp-<br>**PDBTitle:** crystal structures of collagen model peptides (pro-pro-gly)2 4-hyp-hyp-gly-(pro-pro-gly)4 |
| 35 | c3a08E_ | Alignment | not modelled | 18.0 | 58 | **PDB header:**structural protein<br>**Chain:** E: **PDB Molecule:**collagen-like peptide;<br>**PDBTitle:** structure of (ppg)4-oog-(ppg)4, monoclinic, twinned crystal |
| 36 | c3a1hA_ | Alignment | not modelled | 17.6 | 60 | **PDB header:**structural protein<br>**Chain:** A: **PDB Molecule:**collagen-like peptide;<br>**PDBTitle:** crystal structure analysis of the collagen-like peptide, (ppg)4-otg-2 (ppg)4 |
| 37 | c3a19C_ | Alignment | not modelled | 17.6 | 57 | **PDB header:**structural protein<br>**Chain:** C: **PDB Molecule:**collagen-like peptide;<br>**PDBTitle:** structure of (ppg)4-oog-(ppg)4_h monoclinic, twinned crystal |
| 38 | c3a19E_ | Alignment | not modelled | 17.6 | 57 | **PDB header:**structural protein<br>**Chain:** E: **PDB Molecule:**collagen-like peptide;<br>**PDBTitle:** structure of (ppg)4-oog-(ppg)4_h monoclinic, twinned crystal |
| 39 | c2bf9A_ | Alignment | not modelled | 17.2 | 21 | **PDB header:**hormone<br>**Chain:** A: **PDB Molecule:**pancreatic hormone;<br>**PDBTitle:** anisotropic refinement of avian (turkey) pancreatic polypeptide at 0.2 99 angstroms resolution. |
| 40 | c2dezA_ | Alignment | not modelled | 16.8 | 32 | **PDB header:**neuropeptide<br>**Chain:** A: **PDB Molecule:**peptide yy;<br>**PDBTitle:** structure of human pyy |
| 41 | c3a0mB_ | Alignment | not modelled | 16.1 | 60 | **PDB header:**structural protein<br>**Chain:** B: **PDB Molecule:**collagen-like peptide;<br>**PDBTitle:** structure of (ppg)4-ovg-(ppg)4, monoclinic, twinned crystal |
| 42 | c3a0mD_ | Alignment | not modelled | 16.1 | 60 | **PDB header:**structural protein<br>**Chain:** D: **PDB Molecule:**collagen-like peptide;<br>**PDBTitle:** structure of (ppg)4-ovg-(ppg)4, monoclinic, twinned crystal |
| 43 | c5kc5A_ | Alignment | not modelled | 15.9 | 0 | **PDB header:**signaling protein<br>**Chain:** A: **PDB Molecule:**cerebellin-1;<br>**PDBTitle:** crystal structure of the cbln1 c1q domain trimer |
| 44 | c1x1kF_ | Alignment | not modelled | 15.9 | 57 | **PDB header:**structural protein<br>**Chain:** F: **PDB Molecule:**host-guest peptide (pro-pro-gly)4-(pro-allohyp-<br>**PDBTitle:** host-guest peptide (pro-pro-gly)4-(pro-allohyp-gly)-(pro-2 pro-gly)4 |
| 45 | c3a0mC_ | Alignment | not modelled | 15.1 | 60 | **PDB header:**structural protein<br>**Chain:** C: **PDB Molecule:**collagen-like peptide;<br>**PDBTitle:** structure of (ppg)4-ovg-(ppg)4, monoclinic, twinned crystal |
| 46 | d1pk6c_ | Alignment | not modelled | 14.2 | 18 | **Fold:**TNF-like<br>**Superfamily:**TNF-like<br>**Family:**TNF-like |
| 47 | c3a1hE_ | Alignment | not modelled | 14.2 | 60 | **PDB header:**structural protein<br>**Chain:** E: **PDB Molecule:**collagen-like peptide;<br>**PDBTitle:** crystal structure analysis of the collagen-like peptide, (ppg)4-otg-2 (ppg)4 |
| 48 | c3a1hD_ | Alignment | not modelled | 14.2 | 60 | **PDB header:**structural protein<br>**Chain:** D: **PDB Molecule:**collagen-like peptide;<br>**PDBTitle:** crystal structure analysis of the collagen-like peptide, (ppg)4-otg-2 (ppg)4 |
| 49 | c3a1hC_ | Alignment | not modelled | 14.2 | 60 | **PDB header:**structural protein<br>**Chain:** C: **PDB Molecule:**collagen-like peptide;<br>**PDBTitle:** crystal structure analysis of the collagen-like peptide, (ppg)4-otg-2 (ppg)4 |
| 50 | c1ronA_ | Alignment | not modelled | 14.1 | 26 | **PDB header:**neuropeptide<br>**Chain:** A: **PDB Molecule:**neuropeptide y;<br>**PDBTitle:** nmr solution structure of human neuropeptide y |
| 51 | c2ka3C_ | Alignment | not modelled | 13.7 | 27 | **PDB header:**structural protein<br>**Chain:** C: **PDB Molecule:**emilin-1;<br>**PDBTitle:** structure of emilin-1 c1q-like domain |
| 52 | c4oumA_ | Alignment | not modelled | 13.5 | 15 | **PDB header:**signaling protein<br>**Chain:** A: **PDB Molecule:**caprin-2;<br>**PDBTitle:** crystal structure of human caprin-2 c1q domain |
| 53 | c3a1hF_ | Alignment | not modelled | 13.1 | 59 | **PDB header:**structural protein<br>**Chain:** F: **PDB Molecule:**collagen-like peptide;<br>**PDBTitle:** crystal structure analysis of the collagen-like peptide, (ppg)4-otg-2 (ppg)4 |
| 54 | c3a0mE_ | Alignment | not modelled | 13.1 | 60 | **PDB header:**structural protein<br>**Chain:** E: **PDB Molecule:**collagen-like peptide; |

| | | | | | | |
|---|---|---|---|---|---|---|
| 54 | c3a0mE_ | Alignment | not modelled | 13.1 | 60 | **PDBTitle:** structure of (ppg)4-ovg-(ppg)4, monoclinic, twinned crystal |
| 55 | c4by2C_ | Alignment | not modelled | 13.0 | 9 | **PDB header:**structural protein<br>**Chain:** C: **PDB Molecule:**anastral spindle 2, sas 4;<br>**PDBTitle:** sas-4 (dcpap) tcp domain in complex with a proline rich motif of ana22 (dstil) of drosophila melanogaster |
| 56 | c3ah9F_ | Alignment | not modelled | 12.8 | 57 | **PDB header:**structural protein<br>**Chain:** F: **PDB Molecule:**collagen-like peptide;<br>**PDBTitle:** crystal structure of (pro-pro-gly)9 at 1.1 a resolution |
| 57 | c3ah9A_ | Alignment | not modelled | 12.8 | 57 | **PDB header:**structural protein<br>**Chain:** A: **PDB Molecule:**collagen-like peptide;<br>**PDBTitle:** crystal structure of (pro-pro-gly)9 at 1.1 a resolution |
| 58 | c3admC_ | Alignment | not modelled | 12.5 | 60 | **PDB header:**structural protein<br>**Chain:** C: **PDB Molecule:**collagen-like peptide;<br>**PDBTitle:** crystal structure of (pro-pro-gly)4-hyp-ser-gly-(pro-pro-gly)4 |
| 59 | d1pk6a_ | Alignment | not modelled | 12.0 | 36 | **Fold:**TNF-like<br>**Superfamily:**TNF-like<br>**Family:**TNF-like |
| 60 | d1pk6b_ | Alignment | not modelled | 11.9 | 27 | **Fold:**TNF-like<br>**Superfamily:**TNF-like<br>**Family:**TNF-like |
| 61 | c3a1hB_ | Alignment | not modelled | 11.4 | 60 | **PDB header:**structural protein<br>**Chain:** B: **PDB Molecule:**collagen-like peptide;<br>**PDBTitle:** crystal structure analysis of the collagen-like peptide, (ppg)4-otg-2 (ppg)4 |
| 62 | c3a19F_ | Alignment | not modelled | 11.4 | 52 | **PDB header:**structural protein<br>**Chain:** F: **PDB Molecule:**collagen-like peptide;<br>**PDBTitle:** structure of (ppg)4-oog-(ppg)4_h monoclinic, twinned crystal |
| 63 | c3a08D_ | Alignment | not modelled | 11.4 | 52 | **PDB header:**structural protein<br>**Chain:** D: **PDB Molecule:**collagen-like peptide;<br>**PDBTitle:** structure of (ppg)4-oog-(ppg)4, monoclinic, twinned crystal |
| 64 | c2d3hD_ | Alignment | not modelled | 11.4 | 52 | **PDB header:**structural protein<br>**Chain:** D: **PDB Molecule:**collagen model peptides (pro-pro-gly)4-hyp-hyp-<br>**PDBTitle:** crystal structures of collagen model peptides (pro-pro-gly)2 4-hyp-hyp-gly-(pro-pro-gly)4 |
| 65 | c6a2wA_ | Alignment | not modelled | 11.4 | 27 | **PDB header:**photosynthesis<br>**Chain:** A: **PDB Molecule:**protein fucoxanthin chlorophyl a/c protein;<br>**PDBTitle:** crystal structure of fucoxanthin chlorophyll a/c complex from2 phaeodactylum tricornutum |
| 66 | c6gwxA_ | Alignment | not modelled | 11.3 | 40 | **PDB header:**structural protein<br>**Chain:** A: **PDB Molecule:**optimised ppa-tyr;<br>**PDBTitle:** stabilising and understanding a miniprotein by rational design. |
| 67 | c4jglA_ | Alignment | not modelled | 11.2 | 26 | **PDB header:**structural genomics, unknown function<br>**Chain:** A: **PDB Molecule:**hypothetical protein;<br>**PDBTitle:** crystal structure of a streptavidin-like protein (bacegg_01519) from2 bacteroides eggerthii dsm 20697 at 1.25 a resolution |
| 68 | c4d7yA_ | Alignment | not modelled | 11.1 | 23 | **PDB header:**signaling protein<br>**Chain:** A: **PDB Molecule:**c1q-related factor;<br>**PDBTitle:** crystal structure of mouse c1ql1 globular domain |
| 69 | c3admD_ | Alignment | not modelled | 11.0 | 60 | **PDB header:**structural protein<br>**Chain:** D: **PDB Molecule:**collagen-like peptide;<br>**PDBTitle:** crystal structure of (pro-pro-gly)4-hyp-ser-gly-(pro-pro-gly)4 |
| 70 | c4unmB_ | Alignment | not modelled | 11.0 | 15 | **PDB header:**metal binding protein<br>**Chain:** B: **PDB Molecule:**secreted protein;<br>**PDBTitle:** structure of galactose oxidase homologue from streptomyces lividans |
| 71 | c4r8oB_ | Alignment | not modelled | 10.9 | 11 | **PDB header:**structural genomics, unknown function<br>**Chain:** B: **PDB Molecule:**uncharacterized protein;<br>**PDBTitle:** crystal structure of a duf3836 family protein (bvu_1206) from2 bacteroides vulgatus atcc 8482 at 2.50 a resolution |
| 72 | c3a0mA_ | Alignment | not modelled | 10.8 | 60 | **PDB header:**structural protein<br>**Chain:** A: **PDB Molecule:**collagen-like peptide;<br>**PDBTitle:** structure of (ppg)4-ovg-(ppg)4, monoclinic, twinned crystal |
| 73 | c4r03A_ | Alignment | not modelled | 10.8 | 16 | **PDB header:**structural genomics, unknown function<br>**Chain:** A: **PDB Molecule:**uncharacterized protein;<br>**PDBTitle:** crystal structure of a duf3836 family protein (bdi_3222) from2 parabacteroides distasonis atcc 8503 at 1.50 a resolution |
| 74 | c1o91B_ | Alignment | not modelled | 10.8 | 27 | **PDB header:**structural protein<br>**Chain:** B: **PDB Molecule:**collagen alpha 1(viii) chain;<br>**PDBTitle:** crystal structure of a collagen viii nc1 domain trimer |
| 75 | d1o91a_ | Alignment | not modelled | 10.8 | 27 | **Fold:**TNF-like<br>**Superfamily:**TNF-like<br>**Family:**TNF-like |
| 76 | c3ah9C_ | Alignment | not modelled | 10.5 | 57 | **PDB header:**structural protein<br>**Chain:** C: **PDB Molecule:**collagen-like peptide;<br>**PDBTitle:** crystal structure of (pro-pro-gly)9 at 1.1 a resolution |
| 77 | c3ah9E_ | Alignment | not modelled | 10.5 | 57 | **PDB header:**structural protein<br>**Chain:** E: **PDB Molecule:**collagen-like peptide;<br>**PDBTitle:** crystal structure of (pro-pro-gly)9 at 1.1 a resolution |
| 78 | c3ah9B_ | Alignment | not modelled | 10.5 | 57 | **PDB header:**structural protein<br>**Chain:** B: **PDB Molecule:**collagen-like peptide;<br>**PDBTitle:** crystal structure of (pro-pro-gly)9 at 1.1 a resolution |
| 79 | c3admE_ | Alignment | not modelled | 10.5 | 60 | **PDB header:**structural protein<br>**Chain:** E: **PDB Molecule:**collagen-like peptide;<br>**PDBTitle:** crystal structure of (pro-pro-gly)4-hyp-ser-gly-(pro-pro-gly)4 |
| 80 | c3admF_ | Alignment | not modelled | 10.5 | 60 | **PDB header:**structural protein<br>**Chain:** F: **PDB Molecule:**collagen-like peptide; |

| | | | | | | |
|---|---|---|---|---|---|---|
| 80 | [illegible] | Alignment | not modelled | 10.5 | 60 | **PDBTitle:** crystal structure of (pro-pro-gly)4-hyp-ser-gly-(pro-pro-gly)4 |
| 81 | c3admA_ | Alignment | not modelled | 10.5 | 60 | **PDB header:**structural protein<br>**Chain:** A: **PDB Molecule:**collagen-like peptide;<br>**PDBTitle:** crystal structure of (pro-pro-gly)4-hyp-ser-gly-(pro-pro-gly)4 |
| 82 | c3admB_ | Alignment | not modelled | 10.5 | 60 | **PDB header:**structural protein<br>**Chain:** B: **PDB Molecule:**collagen-like peptide;<br>**PDBTitle:** crystal structure of (pro-pro-gly)4-hyp-ser-gly-(pro-pro-gly)4 |
| 83 | c3mswA_ | Alignment | not modelled | 10.4 | 9 | **PDB header:**structural genomics, unknown function<br>**Chain:** A: **PDB Molecule:**uncharacterized protein;<br>**PDBTitle:** crystal structure of a protein with unknown function (bf3112) from2 bacteroides fragilis nctc 9343 at 1.90 a resolution |
| 84 | c2x12A_ | Alignment | not modelled | 10.4 | 27 | **PDB header:**cell adhesion<br>**Chain:** A: **PDB Molecule:**fimbriae-associated protein fap1;<br>**PDBTitle:** ph-induced modulation of streptococcus parasanguinis adhesion by fap12 fimbriae |
| 85 | c6ieeD_ | Alignment | not modelled | 10.2 | 25 | **PDB header:**motor protein<br>**Chain:** D: **PDB Molecule:**basal-body rod modification protein flgd;<br>**PDBTitle:** crystal structure of the flagellar hook cap from salmonella enterica2 serovar typhimurium |
| 86 | d1c3ha_ | Alignment | not modelled | 10.2 | 23 | **Fold:**TNF-like<br>**Superfamily:**TNF-like<br>**Family:**TNF-like |
| 87 | c5m45K_ | Alignment | not modelled | 10.2 | 13 | **PDB header:**ligase<br>**Chain:** K: **PDB Molecule:**acetone carboxylase beta subunit;<br>**PDBTitle:** structure of acetone carboxylase purified from xanthobacter2 autotrophicus |
| 88 | c1gr3A_ | Alignment | not modelled | 10.1 | 27 | **PDB header:**collagen<br>**Chain:** A: **PDB Molecule:**collagen x;<br>**PDBTitle:** structure of the human collagen x nc1 trimer |
| 89 | d1gr3a_ | Alignment | not modelled | 10.1 | 27 | **Fold:**TNF-like<br>**Superfamily:**TNF-like<br>**Family:**TNF-like |
| 90 | c3a0aE_ | Alignment | not modelled | 10.1 | 60 | **PDB header:**structural protein<br>**Chain:** E: **PDB Molecule:**collagen-like peptide;<br>**PDBTitle:** structure of (ppg)4-opg-(ppg)4, monoclinic, twinned crystal |
| 91 | c3a08B_ | Alignment | not modelled | 10.0 | 55 | **PDB header:**structural protein<br>**Chain:** B: **PDB Molecule:**collagen-like peptide;<br>**PDBTitle:** structure of (ppg)4-oog-(ppg)4, monoclinic, twinned crystal |
| 92 | c3a08A_ | Alignment | not modelled | 10.0 | 55 | **PDB header:**structural protein<br>**Chain:** A: **PDB Molecule:**collagen-like peptide;<br>**PDBTitle:** structure of (ppg)4-oog-(ppg)4, monoclinic, twinned crystal |
| 93 | c3abnC_ | Alignment | not modelled | 9.7 | 60 | **PDB header:**structural protein<br>**Chain:** C: **PDB Molecule:**collagen-like peptide;<br>**PDBTitle:** crystal structure of (pro-pro-gly)4-hyp-asp-gly-(pro-pro-gly)4 at 1.022 a |
| 94 | c6d5fF_ | Alignment | not modelled | 9.5 | 23 | **PDB header:**virus<br>**Chain:** F: **PDB Molecule:**fimbrial protein;<br>**PDBTitle:** cryo-em reconstruction of membrane-enveloped filamentous virus sfv12 (sulfolobus filamentous virus 1) |
| 95 | c2rt4A_ | Alignment | not modelled | 9.5 | 41 | **PDB header:**de novo protein<br>**Chain:** A: **PDB Molecule:**af.2a1;<br>**PDBTitle:** nmr structure of designed protein, af.2a1, (ensembles) |
| 96 | c4f3jA_ | Alignment | not modelled | 9.5 | 23 | **PDB header:**signaling protein<br>**Chain:** A: **PDB Molecule:**complement c1q tumor necrosis factor-related protein 5;<br>**PDBTitle:** crystal structure of trimeric gc1q domain of human c1qtnf5 associated2 with late-onset retinal macular degeneration |
| 97 | c5f1bC_ | Alignment | not modelled | 9.4 | 21 | **PDB header:**viral protein/transport protein<br>**Chain:** C: **PDB Molecule:**niemann-pick c1 protein;<br>**PDBTitle:** structural basis of ebola virus entry: viral glycoprotein bound to its2 endosomal receptor niemann-pick c1 |
| 98 | c3c12A_ | Alignment | not modelled | 9.3 | 15 | **PDB header:**biosynthetic protein<br>**Chain:** A: **PDB Molecule:**flagellar protein;<br>**PDBTitle:** crystal structure of flgd from xanthomonas campestris:2 insights into the hook capping essential for flagellar3 assembly |
| 99 | c3osvC_ | Alignment | not modelled | 9.2 | 35 | **PDB header:**structural protein<br>**Chain:** C: **PDB Molecule:**flagellar basal-body rod modification protein flgd;<br>**PDBTitle:** the crytsal structure of flgd from p. aeruginosa |
